# De novo design of autocatalytically forming intra- and intermolecular isopeptide bonds to construct rigid covalent protein assemblies

**DOI:** 10.64898/2026.08.13.744004

**Authors:** Lukas F. Milles, Evelyn B. Huddy, Ann Carr, Yang Hsia, Xinting Li, Alex Kang, Banumathi Sankaran, Asim K. Bera, David Baker

## Abstract

Isopeptide bonds are amide bonds between amino acid side chains that can form autocatalytically, notably in the pili of Gram-positive bacteria. Here, we design *de novo* proteins that form both intramolecular and intermolecular isopeptide bonds entirely autocatalytically. We report over 50 designs that form isopeptide bonds, validated by mass spectrometry and 5 crystal structures. We redesign these constructs as split proteins that form a covalent intermolecular isopeptide crosslink when combined. These split designs are orthogonal to the existing isopeptide-based SpyTag/Catcher system, and their formation can be regulated by temperature, providing control over the timing of crosslinking in protein assemblies. We extend these designs to create rigid domain crosslinks that enable the construction of large well ordered symmetric rings of up to 215 kDa that are irreversibly covalently crosslinked by multiple isopeptide bonds into a single molecule. Our results provide insight into the determinants of isopeptide bond formation, considerably expand the set of isopeptide bond crosslinking systems, and establish a framework to construct fully covalent rigid protein assemblies.

## Introduction

Control over protein-protein interactions in designed or engineered proteins is typically addressed by optimizing thermodynamic affinities that are ultimately reversible and limited by the dissociation rate of the bound complex(*1–8*). Covalent crosslinks provide virtually “infinite affinity”(*9*, *10*) between proteins allowing the construction of stable higher order assemblies and permit irreversible tagging of proteins. Isopeptide bonds are amide bonds that can form (*11*, *12*) between compatible amino acid side chains, a primary amine on a lysine reacting for example with a carbonyl group on aspartic acid(*13*). Isopeptide bonds can be formed by enzymes, for example ubiquitin ligases or transglutaminases. Distinctly different from their role in protein degradation signalling or extracellular matrix stabilization, isopeptide bonds were discovered (*14*) to be formed autocatalytically through loss of an H_2_O or HN_3_ (for ASP and ASN respectively) of Ig-like stalk and pili domains of Gram-positive bacteria via a buried reactive triad in a hydrophobic environment(*15*) of LYS, ASN or ASP, and a catalytic GLU or very rarely ASP(*16*). These isopeptides stabilize both the intermolecular linkage of pilin subunits that extend a tip adhesin domain and the intramolecular covalent locking of the N- and C-termini of protein domains subjected to external stress. Isopeptide formation protects these proteins against mechanical stress, thermal denaturation and proteolytic digestion(*17*). Many of these isopeptide bonds form autocatalytically when the proteins fold, requiring neither enzymes nor special cofactors. The SpyTag/Catcher family(*18–22*) of covalent tagging systems were derived from splitting isopeptide forming domains into a peptide tag and remaining catcher domain that when combined reconstitute the full protein and form the covalent isopeptide crosslink between them. These systems have become widely adopted, versatile, and reliable covalent protein tagging systems(*23*, *24*). Isopeptide bond forming proteins have been engineered by transplanting the reactive triad into a structurally homologous Ig-like domain (*25*) and more recently by scaffolding and designing structures around the isopeptide active site motif of the C-terminal Rrga Ig-like fold (*26*). However, overall few isopeptide forming proteins have been engineered, and there are no such systems that enable construction of rigid protein assemblies (*27*).

We set out to explore the design of an expansive set of *de novo* proteins that can form intramolecular and intermolecular isopeptide bonds autocatalytically, to create orthogonal, split isopeptide systems. We reasoned that a broad range of new isopeptide forming proteins could be designed by starting from isopeptide motif geometries found in nature, and generating new protein backbones scaffolding these catalytic motifs of these enzymes (*28*) using recently developed deep learning methods (*29–31*) such as deep network-based protein hallucination, protein inpainting (*32*) and RFDiffusion(*33*). We set out to design autocatalytic isopeptide bonds *de novo* using these deep learning design methods (*34*), and to explore the use of the designs to generate rigid, fully covalently crosslinked assemblies.

## Results

### Computational design of isopeptide bond forming de novo proteins

We began by characterizing the conserved isopeptide bond forming and adjacent residues in 60 crystal structures of isopeptide bond containing structures (processed input PDB files in Supplementary Data). The autocatalytic amino acid triad of a Lysine (LYS) reacting with an Asparagine or Aspartic acid (ASN, ASP, both indicated as ASX), catalysed by a Glutamic Acid (GLU) positioned between them, are buried within the core of the protein often covered by bulky hydrophobic residues. LYS and ASX are positioned on the N- and C-terminal paired parallel beta strands of these folds. The ASX is +2 register positions downstream relative to the Lysine position (Fig. 1A).

**Fig. 1.**
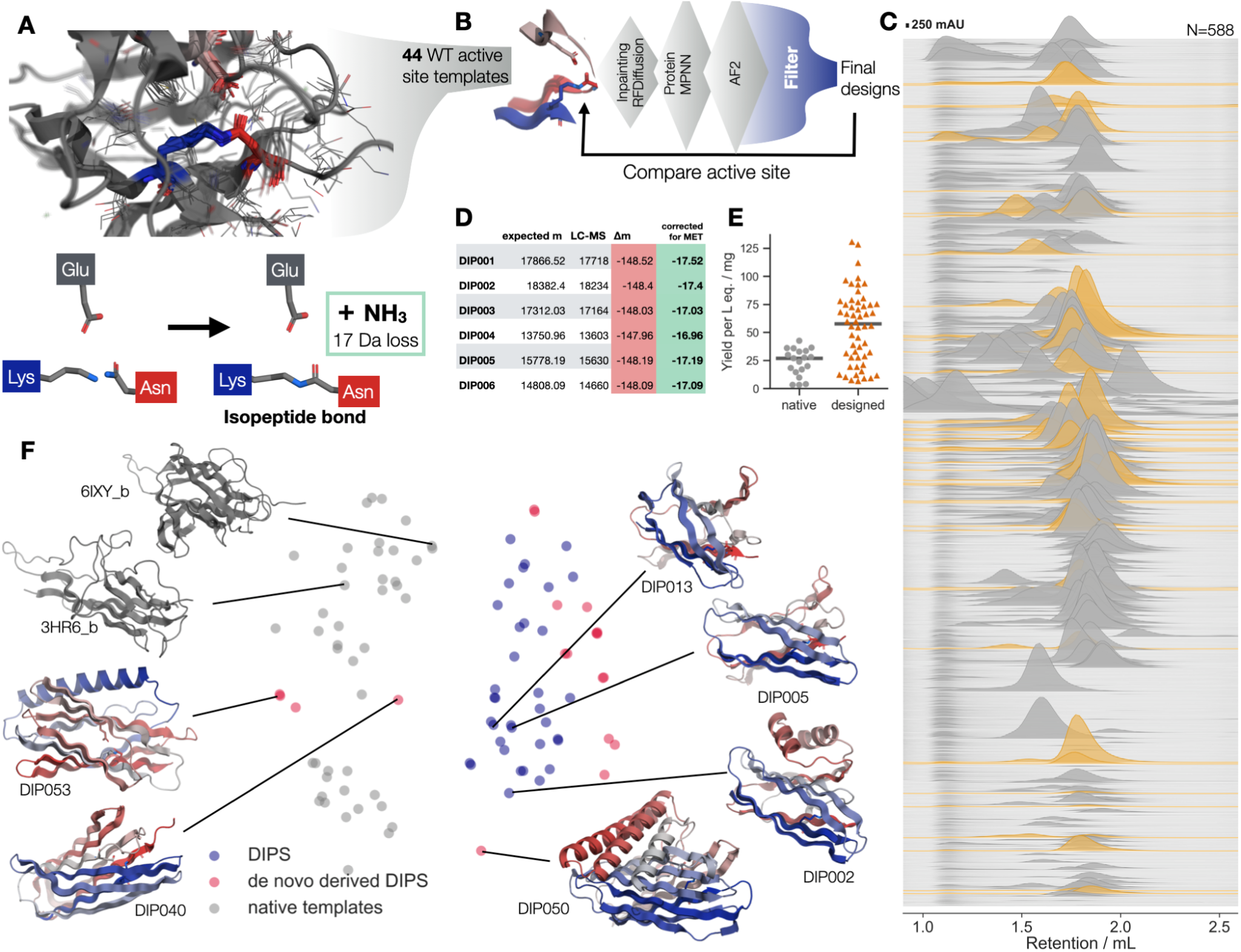
*De novo* design of structurally diverse folds that autocatalytically form intramolecular isopeptide bonds. (A) Isopeptide bond forming reaction LYS + ASN for a side chain amide bond with a leaving NH_3_ group of 17 Da, top - structural alignment of 20 native crystal structures show autocatalytic isopeptide bond formation LYS in Blue, ASX in red, catalytic GLU in pale red (B) Design strategy: structural motifs derived from WT isopeptide domains are used to template protein inpainting or RFDiffusion, backbones are selected and given coding sequences using ProteinMPNN, either manually of explicitly enforcing the reactive triad. AF2 structure predictions are used as the final filter for ordering ensuring that the reactive triad is buried in the core of protein and LYS-ASX are sufficiently close (LYS-NZ to ASX-CG < 4.5 Å, (*53*)) for the isopeptide bond to form (C) Experimental results, 588 SEC chromatograms (Cytiva S75 Increase GL 5/150) in PBS of DIPS and their corresponding retention, isopeptide forming designs are shown in orange, showing clear monodisperse peaks, albeit sometimes void or homodimer peaks too. Active site ALA scan controls showing that with one amino acid of the reactive triad mutated to ALA designs can still be soluble, but do not show the 17 Da mass loss indicative of isopeptide formation can be found in Fig. S8 demonstrating that designs are stable even without the side chain stabilization or “locking” the proteins termini together. (D) LC-MS table showing the mass losses indicative of isopeptide formation for 6 designs (E) Comparison of yield per liter equivalent of 19 native isopeptide forming domains and the 54 isopeptide forming DIPS (F) Structural diversity of DIPS: Multi Dimensional Scaling (MDS) of an all-by-all TM-align of all template isopeptide structures (gray) and validated active DIPS (blue), example structures shown (N to C terminus, blue to red) de novo derived DIPS (red) using the motifs extracted from validated DIPS of the first design rounds.

To generate new proteins harboring this minimal autocatalytic motif, we extracted motifs from native structures that included the reactive triad with 3 amino acids preceding and following the active site residues also, while preserving their N- to C-terminal order (Fig. 1B). We used deep network-based protein hallucination, protein inpainting (*32*) and in subsequent design rounds RFDiffusion to generate backbones that could scaffold these motifs, typically 1000 designs per each of the over 40 templates (Fig. 1B). In the generated protein backbones we confirmed that distances between the C alpha residues constituting the reactive triad were sufficiently close for the side chains to react. Sequences for these backbones were designed using ProteinMPNN(*35*), preserving the identities of the isopeptide forming residues, and optionally also their immediate neighbors. Designs for which the isopeptide forming geometry was recapitulated in the AlphaFold2 (AF2) structure predictions (single sequence mode, no MSA (*36*)), and the designed topology was different from the template structure (TM-score < 0.6) were selected for experimental testing. We refer to these *D*esigned *I*sopeptide bond forming *P*rotein*S* as DIPS throughout this paper.

### Experimental screening, validation, and characterization of isopeptide bond forming designs

Linear DNA fragments encoding DIPS ranging from 90-150 amino acids in length from the first design rounds, were synthesized and cloned into the pre-made background-reducing vector (cwby0001/LM0627) using Golden Gate Assembly (GGA) to append a C-terminal His-tag. Expression and purification were carried out using the SAPP pipeline(*37*) (Supplementary Information). Of 588 designs tested (in four experimental rounds of 82, 192, 120, and 192 respectively), 380 (65%) were soluble and 212 (36%) exhibited a monodisperse peak at or close to the anticipated molecular weight, as determined by Size Exclusion Chromatography (SEC). Compared to 19 positive control WT isopeptide template template structures, the final 54 DIPS (Structures, Fig. S1) had a higher median protein yield of 58 compared to 27 mg per liter equivalent for the native folds (Fig. 1C,E native isopeptide domain SEC in Fig. S2, all DIPS SEC traces in Fig. S3).

To assess isopeptide bond formation, SEC-purified samples were tested by liquid chromatography intact protein mass spectrometry (LC-MS). The designs all have a reactive triad of LYS, ASN with a catalytic GLU; when LYS and ASN react to form the side chain amide bond an NH_3_ group leaves, reducing the total mass of the protein by 17 Da upon isopeptide bond formation. In total 54 of the designs had a peak in the deconvoluted mass spectrum at 148 Da less than the expected molecular weight calculated from the amino acid sequence. Of this mass loss, 131 Da are due to N-terminal methionine cleavage in *E. Coli* (*38*) and the remaining 17 Da thus most likely are the result of isopeptide bond formation (Fig. 1D). Some designs showed partial isopeptide formation as indicated by a second peak with the mass of an unreacted intact protein (for an overview see Supplementary Data table and Fig. S4.). Peak amplitudes in the deconvoluted spectra do not quantitatively reflect relative populations, but qualitatively, after long incubations at 4 °C (∼ 3-4 weeks) the isopeptide bond containing peak grew while the unreacted peak shrunk, indicating that while the reaction in these designs is slow, it is also irreversible as an increase in the non-reacted peak was never observed (Fig. S5). We found that the reactivity of the partially reacting designs could be increased in some cases (Fig. S6) by addition of a large bulky hydrophobic PHE or TRP downstream of the isopeptide bond on the beta strand containing the LYS. DIP001 showed the expected Circular Dichroism (CD) spectrum for an all beta fold, and exhibited no loss of its all beta strand secondary structure up to 95 °C (Fig. S7).

To validate that the isopeptide bonds were in the designed location we tested knockouts for each crosslinking design, individually replacing each reactive amino acid of the triad with an ALA. As expected, these designs - when soluble and monodisperse on SEC - did not show the 17 Da mass loss of isopeptide bond formation (Fig. S8). Although reactive designs with LYS and ASX in swapped positions yielded soluble protein, LC-MS analysis did not show isopeptide bond formation, indicating the importance of the LYS-ASX sequence, which was not initially expected (Fig. S9). The “closing” or “protective” bulky, hydrophobic (PHE, TRP) residue 2 amino acids downstream of the LYS residue was crucial to retain, or in some cases improve activity (Fig. S6). However, a clear “recipe” for bond formation similar to (*25*, *26*) did not emerge.

### Extended design round based on DIPS active sites as templates

Finally, to explore generalizability, we used DIPS as templates for another design round. We extracted isopeptide forming motifs from first design round *de novo* DIPS based on their AF2-predicted models, and scaffolded these motifs with RFDiffusion (*33*) into more *de novo* folds. Experimentally, these were less successful than the first three rounds (11 isopeptide formed out of 192, 6%), with only one design (DIP046) achieving quantitative isopeptide bond yield, though these designs are more varied in structure and motif placement and size. For example, DIP037 with 242 amino acids had an unusual mixed alpha-beta topology and the isopeptide bond was not at the N- and C-termini of the protein. (Fig S3 and Fig. S5). Overall these designs up to 250 aa in length show high diversity and are structurally different as indicated by Multi Dimensional Scaling of an TM-align-based all-by-all alignment when compared to their WT templates (Fig. 1F).

### Structural validation and novelty of designs

To evaluate the accuracy of DIPS designs, we solved 5 crystal structures (Fig 2). These structures show continuous electron density between the isopeptide forming LYS and ASN residues, consistent with formation of the amide bond between the expected side chains. Designs match their experimental structures well with all backbone TM-align RMSDs < 1.5 Å. In one case there is a minor discrepancy between designs and crystal structures, a part of an edge strand of DIP001 predicted to be discontinuous pairing is a straight sheet in the crystal structure. In DIP014 a looped part of a beta strand turn (residue 89-99) is forming a helix instead. DIP046, the only crystal structure of a DIPS motif-based design, shows excellent rotamer agreement and an unusually deep burial of the isopeptide bond, which in native structures is often found at the C-terminal end of paired beta sheets. Taken together, these design efforts result in 54 *de novo* isopeptide forming proteins that are new to nature, with the closest structural homologs detected by foldseek (against the PDB100) (*39*) close to or below to a TM-score (*40*) of 0.5. For example given DIP008 and DIP017 foldseek only finds structural homologs with a TM-score < 0.4 (Fig. S10).

**Fig. 2.**
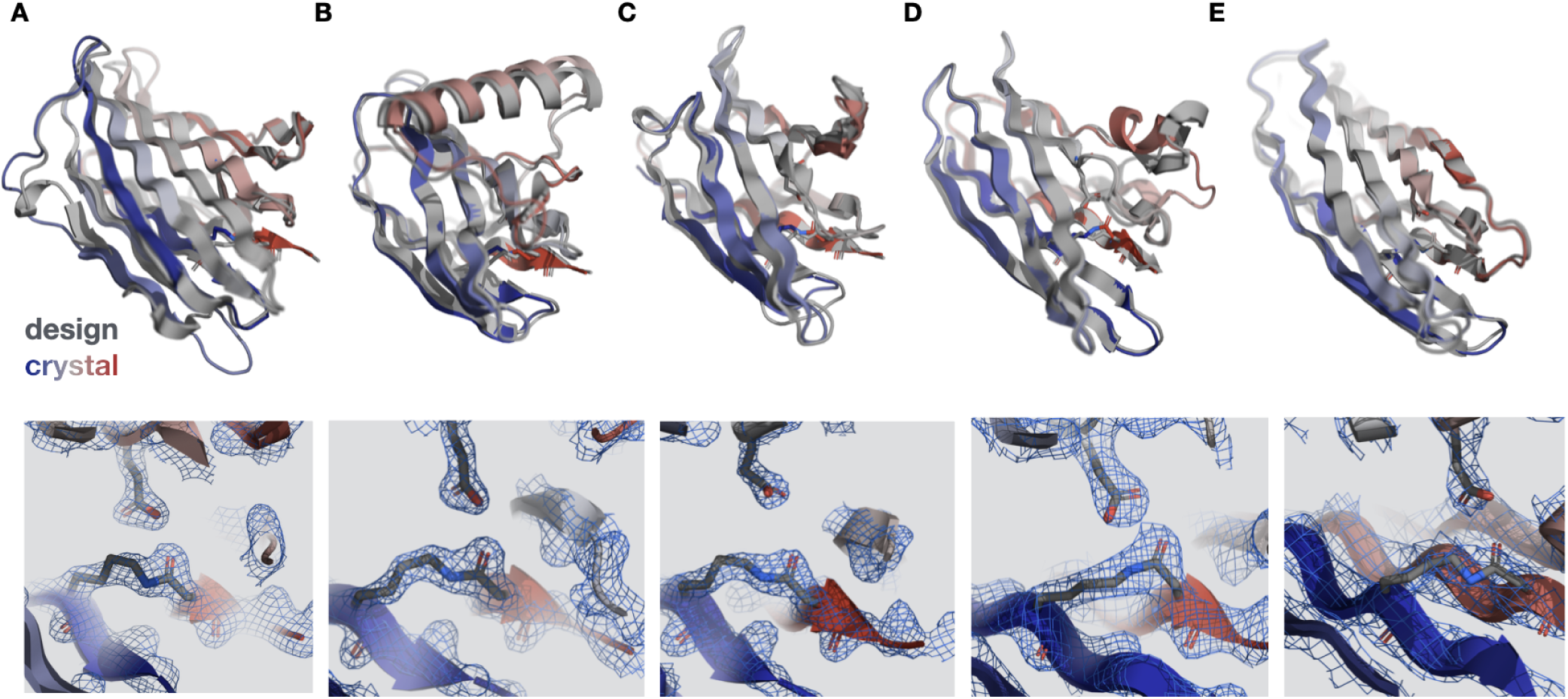
Structural validation of isopeptide bonds formation with 5 crystal structures. For all: top TM-align of crystal structure (blue to red) against design model (gray), bottom 2mFo-DFc map as blue mesh overlaid with the isopeptide bond demonstrating the covalent side chain linkage. TM-align score for each design: (A) DIP001, TM-score: 0.91, RMSD: 1.57 Å, PDB 9N2X, (B) DIP003, TM-score: 0.95, RMSD: 1.04 Å, PDB 9N2Y, (C) DIP013, TM-score: 0.91, RMSD: 1.36 Å, PDB 9N2Z, (D) DIP014, TM-score: 0.92, RMSD: 1.75 Å, PDB 9N30, (E) DIP046, TM-score: 0.96, RMSD: 0.94 Å, PDB 9NMZ, This design’s active site is based on the *de novo* protein DIP019.

### Design and experimental characterization of split isopeptide proteins as covalent crosslinks

To create intermolecular isopeptide crosslinks we split the domains into a peptide and a “catcher” domain, analogously to the Spytag/Catcher system(*19*, *41*). Either the split was performed manually by directly splitting the chain of the DIPS at the N- or C-terminal beta sheet of each domain without any alteration in the sequence. Alternatively, split domains were generated by excising the N- or C-terminal beta strand and these structures were then redesigned using ProteinMPNN(*35*) in a multi-state setup(*42*), jointly optimizing the sequences of the peptide bound complex and peptide unbound, open state (Fig. 3A). In the split-DIPS (sDIPS) complex, the peptide we term DIPtag, the remaining binding domain DIPcatcher. Successful designs were those that in AF2 predictions retained close similarity (< 1.5 A RMSD) to the active site engaged fold of the template PDB, and had a confident (pTM > 0.8) AF2 prediction of the unbound state, ideally with an open cleft for the peptide to bind into. The DIPtag peptide (10-12 amino acids) and DIPcatcher (100-120 amino acids)

**Fig. 3.**
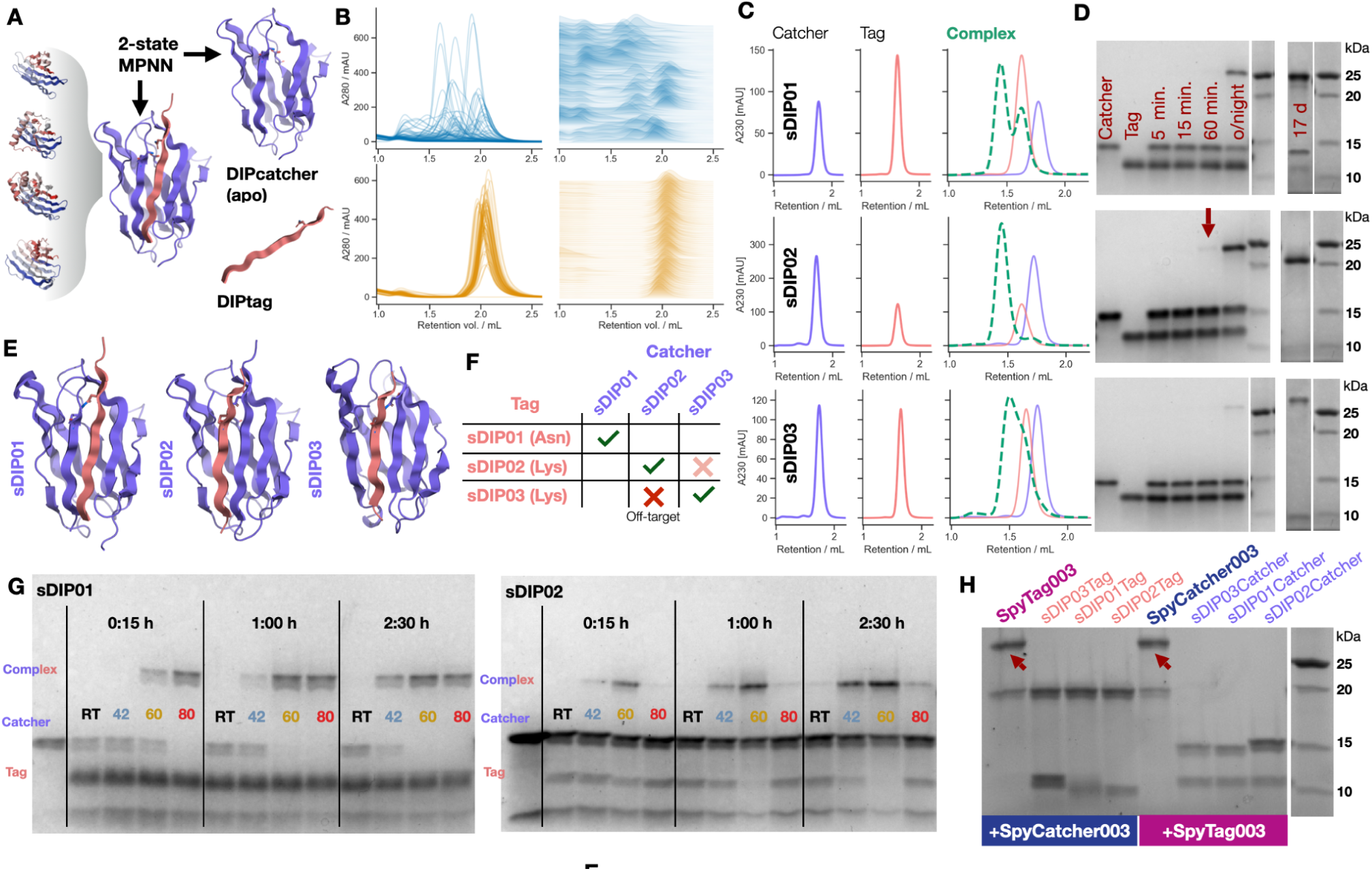
Split-DIPS as Tag/Catcher systems form intermolecular isopeptide bonds slowly but can be accelerated by heating. (A) DIPS were split into DIPTag (peptide, either N- or C-terminal carrying the reactive LYS or ASN, respectively), and DIPCatcher (remaining domain, with open cleft to receive peptide). Muti-state MPNN was used to optimize the apo and peptide bound state. (B) SEC of the catcher designs blue) showing aggregate and void peaks compared to the monodisperse peptides (orange) fused to protein GB1, over half of all catcher domains fail to produce a monodisperse SEC peak. (C) Successful designs showing complex formation between DIPtag and DIPcatcher: monodisperse SEC of Catcher shown in blue, DIPtag with solubility fusion protein in orange. An overlay of SEC traces and the DIPtag/catcher mixture (green, dashed) shows a clear shift to higher molecular weight. When analysing varying incubation times of the higher molecular weight complex elution peak from SEC on SDS-PAGE a faint complex band at the expected molecular weight is barely detectable after 1h (red arrow), stronger bands only emerge after overnight incubation, indicating that affinity between DIPtag/catcher is high, but the isopeptide bond reaction is slow. After 17 days of incubation the expected almost quantitative yield product (showing terminal digest due to long incubation time) can be seen on SDS-PAGE (E) Models of designs with monodisperse peaks of catcher domains and intermolecular isopeptide bond formation, always as DIPcatcher/DIPtag sDIP01 (C-terminal beta sheet with ASP on tag), sDIP02 (N-terminal beta sheet with LYS on tag), sDIP03 (N-terminal beta sheet with LYS on tag) (F) Confusion matrix of intermolecular bond formation (24 h at 60 °C, SDS-PAGE). There is crossreactivity between compatible LYS carrying tags and catchers, with sDIP02 and sDIP03 showing only faint complex bands. (G) Heating up to 60 °C accelerates intermolecular isopeptide bond formation (here at 10 µM equimolar concentration for the orthogonal pair of sDIP01 and sDIP02), in the case of sDIP01 even at 80 °C isopeptide bonds still form. Incubation at 60 °C for 1 h can be used to produce near quantitative yield for all sDIPS. (H) DIPtag and -Catcher are orthogonal to the SpyTag/Catcher003 isopeptide bond forming system, showing no aberrant complex formation after 2 h at 60 °C. Cognate Spytag/Catcher covalent complexes marked by red arrows.

Experimental testing of the sDIPS was conducted with the split peptide expressed as a fusion to a solubility tag such as redesigned Protein A, SUMO, Protein G or the small dimeric FIVAR domain from C. perfringens (*43*). DIPTag and Catcher were expressed separately, and later combined at equimolar concentrations (10 or 20 µM) to test isopeptide bond formation after defined time intervals. While expression of the DIPtag peptide fused to small proteins had almost no solubility issues 153 of 159 (96%) were soluble and monodisperse, only 42 of 172, (24%) of the DIPcatcher domains were soluble and showed monodisperse elution on SEC (Fig. S11). Most DIPcatchers failed due to insolubility, or formation of higher order oligomers, less than 42 ran as monodisperse peak of expected molecular weight on a calibrated SEC column (Fig. 3B). When DIPcatchers were mixed with their peptide target after at least 1 h incubation these ran at the expected complex elution volume on SEC (Fig. 3C), however when testing those Catcher-tag complex fractions in SDS-PAGE, they showed almost no covalent complex formation - indicating that affinity to the target was strong, but isopeptide bond formation in the complex slow (Fig. 3D).

### Split isopeptide crosslink formation is accelerated by heat and orthogonal to SpyTag/Catcher

Three sDIPS designs showed slow formation of the intermolecular covalent isopeptide bond after 17h of incubation (Fig. 3D,E) at equimolar concentration of 10 µM on SDS-PAGE. We also confirmed by LC-MS the presence of a covalently linked Tag-Catcher complex that was 17 Da lighter than expected loss due to NH_3_ loss from isopeptide bond formation (Fig. S12). Leaving the reaction to continue for approximately 1 month at 4 °C increased product formation to quantitative yields, indicating that the reaction indeed proceeded slowly. Improving these designs by adding additional hydrophobic residues (PHE, TRP) near the active site lead to recurring aggregation, and especially homodimer formation - albeit the homodimers readily reacting with their cognate DIPtag targets. When adding these hydrophobic residues to the DIPtags, especially downstream of the reactive LYS in one case reaction rates improved.

We reasoned that heat would increase isopeptide reaction rates and tested bond formation of split-DIPS at room temperature (RT), 42, 60 and 80 °C. Increasing temperature increased isopeptide product formation when comparing identical incubation times. For two constructs near quantitative bond formation was achieved after only 60 minutes at 60 °C (Fig. 3G).

Two of the DIPtag/catcher constructs cross react to another tag or catcher (Fig. 3F, Fig. S13), as they have compatible N-terminal extracted tag peptides carrying Lysines.DIPtag/catcher systems are orthogonal to the previously developed SpyTag003 / SpyCatcher003 system in SDS-PAGE cross reaction assays, even at elevated temperatures there was no product formation between SpyTag003 or catcher and the split-DIPS visible (Fig. 3F,H). Overall sDIP01, sDIP02 and sDIP03 splitDIPS pairs can be used orthogonally to the SpyTag/Catcher003 system. Other compatible DIPCatcher/Tag systems were functionally validated, though these have in part slower reaction rates or present issues of Catcher aggregation (Fig. S14).

### Design of rigid covalent assemblies based on split isopeptide designs

Protein design has been used to systematically generate a wide range of modular and rigid molecular assemblies(*4*, *27*, *44–48*). An exciting application of intermolecular isopeptide bonds is to enable the design of extremely stable assemblies - similar to the HK97 bacteriophage’s capsid subunits that are isopeptide bond-linked(*49*). In these assemblies all subunits are covalently linked to each other through spontaneously forming isopeptide bonds, creating one large molecule. Using the DIPtag/Catcher this can be achieved more precisely and with more site-specific control compared to unspecific chemical crosslinking. For precisely designing such systems, it is desirable for the covalent bond forming domain(s) to be rigidly connected to the remainder of the designed system with very little flexibility, so that the 3D architecture of the assembly does not vary or flex. The standard SpyCatcher systems do not currently enable such rigid assembly construction easily due to their purely beta sheet structures, which are incompatible with commonly used rigid helical spacer domains(*45*).

To enable construction of such rigid, fully covalently-linked assemblies, small helical protein domains were docked asymmetrically against both the catcher and tag sides of the Split-DIPS in two independent steps using RPXDock(*50*). Docks were filtered for proximity of joinable termini between helical bundle and Split-DIPS chains, and then short loops were diffused between the two structures with RFDiffusion(*33*). The cores of these rigid split-DIPS (rsDIPS) beta sheet complexes were conserved, with all other residue identities allowed to redesign. Buttressed designs predicted to adopt the target structure with AlphaFold2 were tested as heterodimers (Fig 4A-C), they were found to form intermolecular isopeptide bonds confirmed by SDS-PAGE and LC-MS with the characteristic NH_3_ mass loss of 17 Da for the complex (Fig. S15).

**Fig. 4.**
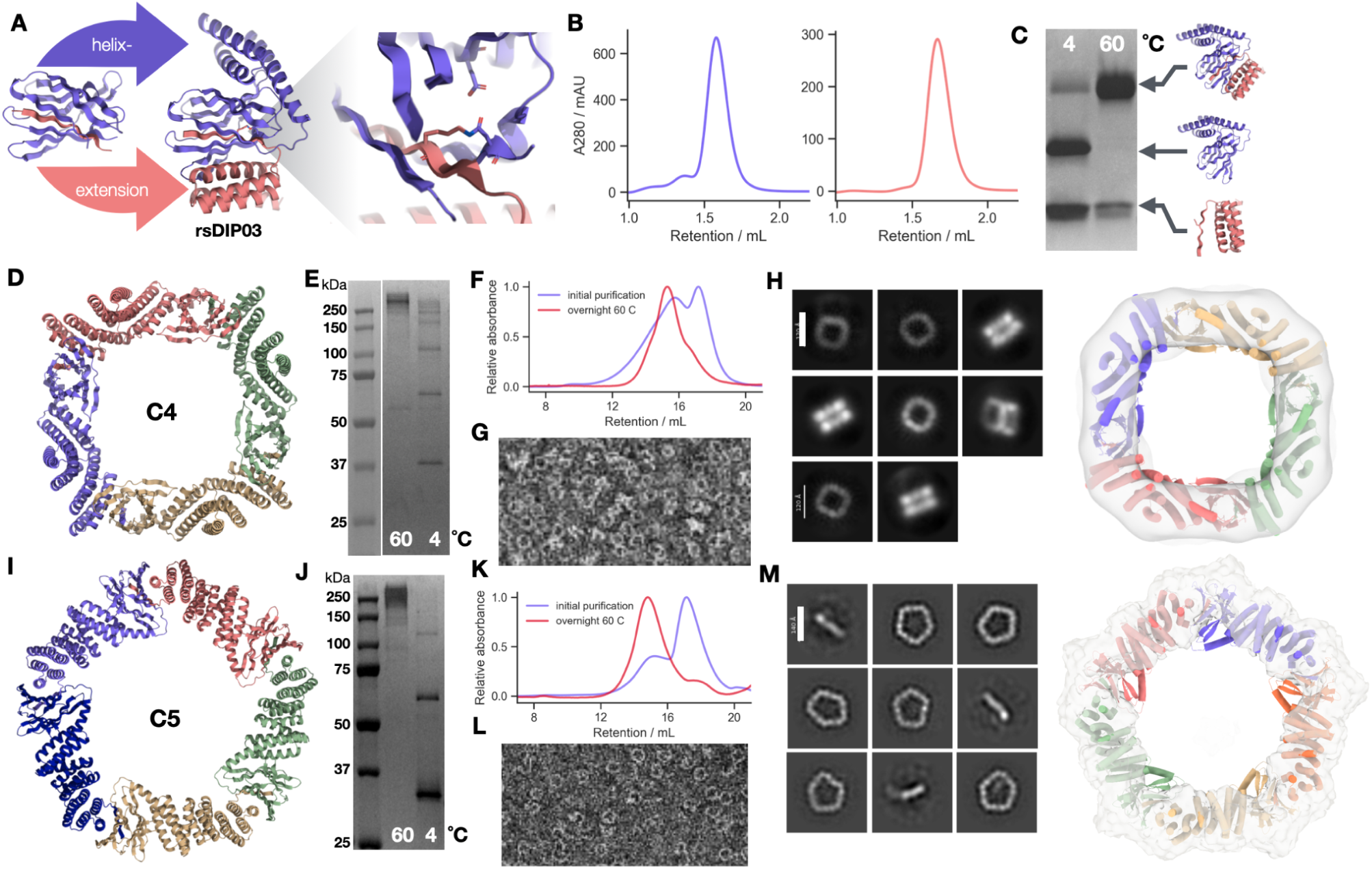
Rigid split-DIPS interfaces allow the construction of fully isopeptide linked oligomeric assemblies. (A) DIPtag and catchers were extended with N- and C-terminal helices using RFDiffusion, the isopeptide forming DIPtag/catcher complex sDIP03 was extended into design rsDIP03, expected isopeptide crosslink between domains shown in AF2 prediction model inset. (B) SEC showing monodisperse peaks of both binding partners in isolation. (C) SDS-PAGE after overnight incubation at 4 and 60 °C showing covalent complex formation (color coded as in structure rendering), isopeptide formation confirmed by LC-MS (Fig. S15) rsDIP03 was designed into a cyclic C4 (D) or C5 (I) homooligomer, where protomer interfaces are linked by isopeptide bonds. (E,J) Purified proteins after overnight incubation at 4 and 60 °C on SDS-PAGE. Heat accelerated isopeptide bond formation shows clear bands at over 150 kDa for both assemblies, in line with the approximate expected molecular weight of C4 (protomer: 40 kDa, oligomer: 161 kDa) C5 (protomer 43 kDa, oligomer 215 kDa). At 4 °C partially reacted oligomers e.g. dimers can be observed. (F,K) qualitative comparison of SEC of the overnight 60 °C heated assemblies (red) compared to the purified sample from expression culture (blue) on Cytiva Superdex S200 shows clear shift to the expected high molecular weight of the C4/C5 rings and only few smaller peaks of likely isopeptide dimers or unreacted protomers after heat annealing. (G,L) Negative stain EM shows open and closed rings formed on the grid, (H,M) class averages confirm the formation of the expected C4 square or C5 pentagon structure, likely fully covalently linked by isopeptide bonds.

To test the ability of Split-DIPS to be used as integral structural elements of protein assemblies, we constructed cyclically symmetrical ring designs which utilize buttressed Split-DIPS as the cyclic interface, in particular, a buttressed design *rsDIP03*, derived from sDIP03 was used. To accomplish this, the WORMS software (*27*) was used to fuse the buttressed catcher on one terminus, a helical adapter protein in the middle, and a buttressed tag on the other terminus such that the overall structure should form a cyclic oligomer upon covalent bond formation. For design C4 derived from rsDIP03, a fourfold cyclically symmetric homoligomer with a square shape formed as designed (Fig. 4 D-H) of 161 kDa; design C5 is a pentagon constructed using the same crosslinking building block rsDIP03 (Fig. 4I-M) of 215 kDa. Heating the protein (60 °C overnight) facilitates complete isopeptide bond formation, leading to a single expected high molecular weight band in SDS-PAGE, showing that temperature drives covalently assembly (Fig 4F,K). Negative-stain EM of the heated sample also shows primarily the expected square C4 and pentagonal C5 species in good agreement with the design models (Fig 4G,L,H,M). Thus, buttressed Split-DIPS enable the formation of rigid fully covalent assemblies with heat induced crosslinks and thus a heat-stable overall fold that is in the end linked into a single molecule.

## Discussion

We design and experimentally confirm *de novo* designed isopeptide bond forming proteins (DIPS) that can be split into heterodimers that form intermolecular isopeptide bonds (split DIPS, sDIPS). These systems can be used to construct rigid fully covalently crosslinked oligomeric assemblies. Our 54 newly designed isopeptide bond forming proteins considerably extend the around 50 crystallized native isopeptide domains resolved in the PDB, providing an opportunity to study isopeptide bond formation free from evolutionary constraints with an extended dataset. The designs presented here could be further refined using Deep Mutational Scanning, directed evolution, and/or active learning approaches to generate for example more orthogonal split DIPS systems. The overall success rates of isopeptide generation - even in later redesign rounds – was low (6-15% of all designs ordered depending on the order). Isopeptide bond formation chemistry, not design solubility or folding, was the main problem; the charged amino acid active triad in the core of the protein does not compromise solubility as much as might be expected, slow reactions rates of such engineered systems had been observed before (*19*, *25*, *26*). Controls of active enzymes, in which only one residue of the reactive triad was mutated to Alanine mostly stayed soluble (Out of 27 designs all show at least a small peak, often at reduced yield, at the expected molecular weight on SEC for at least one, and in 10 cases, for all three mutants, Fig. S8), demonstrated that most scaffolds designed can successfully organize these residues in or close to the core of the protein, although the amino acids are charged (LYS, ASN, GLU).

The 54 *de novo* designed isopeptide bond forming proteins here are blueprints for designed stabilization against mechanical, proteolytic, chemical or heat denaturation(11) of a protein fold by isopeptide bonds as an irreversible alternative to disulfide bonds. Furthermore, DIPS are efficient, albeit autocatalytic, enzyme designs that can achieve quantitative product yield as indicated by LC-MS spectra and crystal structures. The DIPtag/catcher systems developed here form intermolecular isopeptide bonds. While they bind strongly to each other as indicated by complex formation on SEC, they do not immediately react to form the crosslinking isopeptide bonds. Heating the complexes to 60 °C directly accelerates isopeptide bond formation. Thus, in these systems covalent crosslinking can be triggered by heat, allowing for time controlled annealing procedures that can use heat as a stimulus. Potentially, more sophisticated designs using sDIPS and rsDIPS under heating and cooling cycles could allow covalent heterooligomeric assemblies, as binding (at low temperature) and covalent reaction (at higher temperature) can be separated(21). As demonstrated by our fully covalent C4 and C5 designs, the rigid DIPtag/catcher systems enable the construction of higher order fully covalent materials, opening avenues to highly stable covalent symmetric and asymmetric heterooligomeric protein assemblies, fibers or protein nanoparticles. These could have applications in challenging environments as the covalent isopeptide bond will staple subunits together irreversibly, and will resist mechanical tension for example, until such forces cease and the protein can refold (*51*). in novel biomaterials or as enzyme display platforms in harsh chemical environments. Using temperature to control this process, we can “weld”(*52*) subunits together - possibly even using thermal cycling to slowly anneal a material when assemblies are slow-forming. This could allow stepwise construction of complex assemblies with many protein subunits, providing means to a different class of designed protein materials. In sum, we create a new repertoire of *de novo* designed isopeptide crosslinking systems, opening routes to construct fully covalently linked, ultrastable assemblies from simple protein tagging to massive protein nanoparticles.

## Supplementary Information

Supplementary Information containing Table S1 and Figures S1 to S15 is attached to this manuscript.

## Data availability

Source data, sequences, and design models will be made available on Zenodo upon final publication. Crystallographic datasets have been deposited in the PDB with accession codes: 9N2X (DIP001), 9N2Y (DIP003), 9N2Z (DIP013), 9N30 (DIP014), 9NMZ (DIP046). Key plasmids will be deposited with Addgene.

## Competing interests

LFM, EBH, AC, YH, XL, AK, AKB, and DB are coinventors of an invention regarding the designed proteins presented here submitted by the University of Washington. The remaining authors declare no competing interests.

## Acknowledgements

We thank Luki Goldschmidt and Patrick Vecchiato for computational infrastructure, Kandise VanWormer and Hernan Nunez-Ortega for laboratory support. This work was supported by a Human Frontier Science Program Cross Disciplinary Fellowship (LT000395/2020-C, to L.F.M.), an EMBO Non-Stipendiary Fellowship (ALTF 1047-2019, to L.F.M.). This work is based upon research conducted at the Northeastern Collaborative Access Team beamlines, which are funded by the National Institute of General Medical Sciences from the National Institutes of Health (P30GM124165). This research used resources of the Advanced Photon Source, a U.S. Department of Energy (DOE) Office of Science User Facility operated for the DOE Office of Science by Argonne National Laboratory under Contract No. DE-AC02-06CH11357. We also want to thank the Advanced Light Source (ALS) beamline 8.2.1 at Lawrence Berkeley National Laboratory for X-ray crystallography data collection. The Berkeley Center for Structural Biology is supported in part by the National Institutes of Health (NIH), National Institute of General Medical Sciences, and the Howard Hughes Medical Institute. The ALS is supported by the Director, Office of Science, Office of Basic Energy Sciences and US Department of Energy (DOE) (DE-AC02-05CH11231). This research also used AMX of the National Synchrotron Light Source II, a U.S. Department of Energy (DOE) Office of Science User Facility operated for the DOE Office of Science by Brookhaven National Laboratory under Contract No. DE-SC0012704. The Center for BioMolecular Structure (CBMS) is primarily supported by the National Institutes of Health, National Institute of General Medical Sciences (NIGMS) through a Center Core P30 Grant (P30GM133893), and by the DOE Office of Biological and Environmental Research (KP1605010).

## Methods

### Computational design

*De novo* isopeptide bonds were designed using Protein Inpainting (*32*) using the motifs extracted from over 50 template isopeptide domains in the PDB, see Supplementary Data tables. These were centered around their catalytic residues (LYS, ASN/ASP, catalytic GLU). For each WT isopeptide domain motifs were padded with up to 3 neighboring residues and then submitted for inpainting with variable total length of the proteins up to 150 residues. Contigs to connect the motif fragments varied up to the total allowed length of the protein up to 200 amino acids. The order of contigs was fixed from N- to C-terminus LYS, catalytic GLU, ASN/ASP. Backbones were filtered by novelty (TM-score to parent scaffolds < 0.5) and fidelity to the active triad orientation (distance of alpha carbons), as well as other plausibility metrics (DSSP distribution, loop length).

The backbones were then either entirely sequence designed with ProteinMPNN and the reactive triad re-threaded into the correct positions, or the active triad was protected from mutation and only the rest of the protein designed by ProteinMPNN(24). AlphaFold2 (AF2) in single sequence prediction mode (no MSA) was used to assess the MPNN sequence. Designs were filtered mainly by AF2 quality metrics (pTM score > 0.7) and quality of the reconstructed LYS-GLU-ASP active site. Here the tip atom distances between reacting LYS-ASP, and LYS-GLU and LYS-ASP (< 4.2 A). Furthermore the SASA (Solvent Accessible Surface Area) of the reacting tip atoms had to be 0.0 A for the designs to pass filters.

*De novo* designed isopeptide backbones based on motifs now extracted from successful DIPS were designed using RFDiffusion (up to 250 residues) analogously to the strategy employed with protein inpainting, resulting in *de novo* structure derived DIPS.

Isopeptide bonds were into PDB files using Pyrosetta(*54–56*) with modified LYS and ASN residues forming the covalent bond, scripts provided in supplementary data (*57*).

### Cloning

Cloning expression and purification follow the SAPP procedures we developed previously (*37*), software can be downloaded here: https://github.com/bwicky/SAPP_DMX

Golden date entry vectors were propagated and produced from *ccdB*-resistant NEB Stable *E. coli* chemically competent cells (NEB). Following transformation according to the manufacturer’s protocol, the transformants were grown overnight at 37 °C on LB-Agar containing 50 μg/mL of kanamycin sulfate. Single colonies are used to inoculate 50 mL of liquid LB cultures (250 mL baffled conical flasks) with 50 μg/mL of kanamycin sulfate, and grown overnight at 37 °C with shaking at 250 rpm. Entry Plasmids were purified using a ZymoPURE II Plasmid Midiprep Kit (Zymo Research) according to the manufacturer’s protocol.

GGA reactions are assembled with an ECHO 525 acoustic liquid handling robot (Beckman Coulter) by using a custom transfer protocol to assemble the following ratios. A master mix containing BsaI-HFv2 (NEB), T4 DNA ligase (NEB), T4 DNA ligase buffer (NEB) and the entry vector is prepared first, and the ECHO is used to combine master mix and DNA fragments into 96-well PCR plates (1 μL final volumes), here we mostly used cwby001/LM0627 (https://www.addgene.org/191551/) as target vector. Reactions are incubated at 37 °C for at least 20 minutes. GGA products are directly transformed into an *E. coli* expression strain by adding 6 μL of BL21(DE3) chemically competent cells (NEB) to the 1 μL reactions. Transformations are performed in 96-well PCR plates, and the transformed cells are used to inoculate expression cultures directly(*37*).

### Protein expression, purification, SEC, and SDS-PAGE

For screening experiments, expression was carried out according to the SAPP protocol, for a full description refer to(*37*). In brief: Protein expression was performed in round bottom 96 deepwell plates filled with 1 mL of simplified auto-induction media (*58*) (TB II, MP Biomedicals, supplemented with glycerol [5 g/L], glucose [0.5 g/L], lactose [2 g/L], MgSO_4_ [2 mM], and kanamycin sulfate [50 μg/mL]). Transformation reactions were split 4-fold and used to inoculate 4 x 1 mL of expression cultures directly, followed by incubation at 37 °C for *at least* 22 hours at 1,000 rpm.

Cells were harvested in 96 deepwell plates by centrifugation (4,000 x g, 5 minutes), and cells resuspended in lysis buffer (B-PER, Thermo Fisher Scientific, supplemented with lysozyme [0.1 mg/mL], PMSF [1 mM], Benzonase [25 U/mL], 100 μL per 1 mL of culture-equivalent pellet). This chemical lysis ran for15 minutes at 37 °C under agitation (1,000 rpm) before pooling the lysates (4 x 100 μL) into one 96 deep well plate and clearing debris by centrifugation (4,000 x g, 20-30 minutes). Proteins were purified from the soluble fraction by binding to Ni-NTA resin (50 μL of resin bed per well, added to 96-well fritted plates (25 µm PE frit, Agilent Technologies)), followed by three wash cycles using a plate vacuum manifold (3 x 500 μL, 20 mM Tris, 300 mM NaCl, 25 mM imidazole, pH 8.0). Proteins were eluted from the resin by addition of 200 μL of elution buffer (20 mM Tris, 300 mM NaCl, 500 mM imidazole, pH 8.0), and sterile-filtered by centrifugation into a 96-well filter plate (0.22 μm pore-size, Agilent Technologies, 200953) mounted on a receiver plate. Filter-sterilized eluates are further purified by size-exclusion chromatography (SEC) into Phosphate Buffer (PBS) on either a Cytiva Superdex 75 Increase 5/150 GL or Cytiva Superdex 200 Increase 5/150 GL by injecting 100 μL at a flow-rate of 0.65 mL / minutes. Chromatography was performed either on a Agilent 1260 Infinity II Bio-inert system equipped with multisampler and fraction collector (fractionation into 384-well plates), or a Cytvia ӒKTA Pure system equipped with an autosampler and using a modified flow-path. Columns were calibrated using Dextran Blue for void determination and size standards (Cytiva LMW and HMWkits). Chromatograms were batch-analyzed by a custom python script (see github link above) which also generated instructions for an OT-2 liquid handling robot (Opentrons) for cherry-picking, pooling and normalizing fractions for further experiments.

For X-ray crystallography 4x 50 mL cultures in the same Autoinduction media as above were grown for 24h at 37 °C, lysed by sonication, and then purified analogously in 1mL Ni-NTA resin (Thermo Fisher) in a 24 well fritted plate (Agilent Technologies 201415) using appropriately larger Volumes (4 x 10 mL of wash buffer as above). After the the HIS wash buffer step, the resin was exchanged into SNAC Buffer (0.1M CHES-NaOH pH 8.2, 0.1M NaCl, 0.1 M acetone oxime, 0.5 M guanidinium HCl and 2 mM NiCl2)(*59*). The plate was sealed and the suspended resin was incubated in this buffer shaking overnight. Then, after SNAC-tag cleavage, the tagless protein was collected with the flow through. All proteins were then further purified by size exclusion chromatography (in Tris 20 mM, NaCl 50 mM, pH 8) before being concentrated for crystallography (Amicon system, 3 kDa MWCO, Merck Millipore).

All SDS-PAGE gels were stained using coomassie blue (BioRad, TGX Stain free any kD 26 well). Protein samples were diluted into 2x Laemmli Buffer and heated to 95 °C for 10 minutes at 10-20 µM concentrations; gels were run for 20-25 min at 250 constant Voltage.

### Design and assembly of large rings

To make split dips amenable to rigid fusion, they were buttressed with helical bundles on both sides of the split. For each side, a library of helical protein backbones from previously published design work were docked against the split dip asymmetrically with RPXDock(*45*, *50*, *60*). RosettaScripts(*61*) was used to filter for less than 12 A distance of termini of dip chains to termini of helical bundles, and then short linkers were built with RFdiffusion(*33*). Sequences were designed and filtered analogously to the parent DIPS designs.

To generate the cyclic ring structures from these, the WORMS protocol was used as described in the “Crown” structure method in the reference publication(*62*). Inputs to worms were the buttressed split dips and a library of helical protein filler backbones similar to the one used for bundle docking to make the buttresses. Fusion solutions between the two sets of blocks were found that satisfied geometric closure criteria with split dips interfaces acting as cyclic assembly interfaces.

### Mass spectrometry

Intact mass spectra were obtained by reverse-phase LC/MS on an Agilent G6230B TOF using an AdvanceBio RP-Desalting column with a fast gradient of 90:10 (A:B) to 5:95 (A:B) over 2 minutes (A: H2O with 0.1% Formic Acid, B: Acetonitrile with 0.1% Formic Acid), and subsequently deconvoluted with Bioconfirm using a total entropy algorithm from m/z 600-2500 with a fixed mass range between 5000-30000Da. For long time series (days or weeks), samples were incubated in phosphate buffer (PBS) at 4 °C and then re-measured.

### Circular Dichroism

Circular Dichroism (CD) was performed on a Jasco 1500 CD spectrometer with a 6 sample rotating turret. Samples were placed in 1 mm pathlength cuvettes (Hellma QS Quartz cell) at concentrations of 0.2 mg mL^-1^ in PBS (pH 7.4 buffer). The temperature was increased from 25 °C to 95 °C, recording full CD spectra between 200 and 260 nm in 10 °C intervals, and reading at 222 nm in 2 °C intervals. After reaching 95 °C the samples were allowed to cool back to 25 °C before recording a final spectrum.

### X-ray crystallography

All crystallization experiments were conducted using the sitting drop vapor diffusion method. Crystallization trials were set up in 200 nL drops using the 96-well plate format at 20 °C. Crystallization plates were set up using a Mosquito LCP from SPT Labtech, then imaged using UVEX microscopes and UVEX PS-256 from JAN Scientific. Diffraction quality crystals formed in 0.2 M Magnesium chloride, 0.1 M TRIS pH 8.5, and 3.4 M 1,6-Hexanediol for DIP001, in 0.01 M Zinc sulfate, 0.1 M MES pH 6.5, 25 % (v/v) PEG 550 MME for DIP003, in 0.2 M Magnesium chloride hexahydrate, 0.1 M Sodium HEPES pH 7.5, and 30 % (v/v) PEG 400, in for DIP013, in 0.05 M Calcium chloride, 0.1 M BisTris pH 6.5, 30 % (v/v) PEG 550 MME for DIP014 and 0.2 M ammonium citrate dibasic, 20% w/v PEG3350 for DIP046.

Diffraction data was collected either at the Advanced Photon Source beamline on 24-ID-C, Advanced Light Source on Beamline 8.2.1 or NSLS-II beamline 17-ID-1. X-ray intensities and data reduction were evaluated and integrated using XDS(*63*) and merged/scaled using Pointless/Aimless in the CCP4 program suite (*64*). Structure determination and refinement starting phases were obtained by molecular replacement using Phaser (*65*) using the designed model for the structures. Following molecular replacement, the models were improved using phenix.autobuild (4); efforts were made to reduce model bias by setting rebuild-in-place to false, and using simulated annealing and prime-and-switch phasing. Structures were refined in Phenix (4). Model building was performed using COOT (5). The final model was evaluated using MolProbity (6). Data collection and refinement statistics are recorded in Supplementary Table 1. Data deposition, atomic coordinates, and structure factors reported in this paper have been deposited in the Protein Data Bank (PDB), http://www.rcsb.org/ with accession code 9N2X, 9N2Y, 9N2Z, 9N30 and 9NMZ.

### Electron microscopy

Samples for negative-stain electron microscopy were typically prepared at 0.1 mg/mL concentration of protein for initial screening in a PBS buffer. 5 μL was applied on glow discharged, carbon-coated 400-mesh copper grids (01844F, TedPella,Inc.), then washed with Milli-Q Water and stained using 0.75% uranyl formate(*66*). Air-dried grids were then imaged on a FEI Talos L120C TEM (FEI Thermo Scientific, Hillsboro, OR) equipped with a 4K × 4K Gatan OneView camera at a magnification of 57,000x and pixel size of 2.47 Å. Micrographs collection was automated using EPU software (FEI Thermo Scientific, Hillsboro, OR) and were imported into cryoSPARC software(*67*). Typically, micrographs were imported with constant CTF and then manual particle picking of approximately 100 particles was done to make templates through 2D classing for automated picking. After full scale templated particle picking, additional 2D classing was done, and selected 2D classes were used for C1 (non-symmetrized) ab initio reconstruction followed by either homogeneous refinement in the designed cyclic symmetry.

