## Supplementary Figures and Table for "De novo design of autocatalytically forming intra- and intermolecular isopeptide bonds to construct rigid covalent protein assemblies"

This file contains:
Table S1
Figures S1 to S15

**Supplementary Table 1: Crystallographic data collection and refinement statistics**

| | **DIP001/LM1814 (PDB ID: 9N2X)** | **DIP003/LM1843 (PDB ID: 9N2Y)** | **DIP013/LM2363 (PDB ID: 9N2Z)** | **DIP014/LM2379 (PDB ID: 9N30)** | **DIP046/LM4150 (PDB ID: 9NMZ)** |
|---|---|---|---|---|---|
| **Data collection** | | | | | |
| Space group | $P\ 2_1$ | $P\ 2_1\ 2_1\ 2_1$ | $P\ 6_3\ 2\ 2$ | $P\ 3_1\ 2\ 1$ | *H 3 2* |
| Cell dimensions | | | | | |
| *a*, *b*, *c* (Å) | 35.76, 108.50, 45.45 | 40.68, 49.42, 60.16 | 60.91, 60.91, 96.60 | 57.74, 57.74, 226.12 | 162.21, 162.21, 197.00 |
| α, β, γ (°) | 90, 112.91, 90 | 90, 90, 90 | 90, 90, 120 | 90, 90, 120 | 90, 90, 120 |
| Resolution (Å) | 41.86 - 1.81 (1.85 - 1.81) | 60.16 - 1.84 (1.88 - 1.84) | 46.30 - 1.95 (2.00 - 1.95) | 50.01 - 2.20 (2.24 - 2.20) | 34.50 - 3.46 (3.68 - 3.46) |
| $R_{merge}$ | 0.039 (0.941) | 0.471 (2.738) | 0.716 (1.471) | 0.170 (2.668) | 0.668 (2.354) |
| $I / \sigma I$ | 9.9 (0.9) | 4.3 (1.2) | 6.9 (2.2) | 5.5 (0.30) | 4.1 (1.2) |
| Completeness (%) | 89.8 (92.7) | 99.6 (94.0) | 100 (100) | 99.99 (100) | 94.60 (100) |
| Redundancy | 2.6 (2.5) | 12.5 (13.2) | 23.8 (24.1) | 16.3 (17.0) | 12.5 (11.3) |
| **Refinement** | | | | | |
| Resolution (Å) | 41.86 - 1.81 (1.88 - 1.81) | 38.17 - 1.84 (2.02 - 1.84) | 46.30 - 1.95 (2.07 - 1.95) | 50.00 - 2.20 (2.30 - 2.20) | 34.50 - 3.46 (3.60 - 3.46) |
| No. reflections | 25915 (2983) | 10968 (2646) | 8241 (1328) | 21938 (1619) | 12355 (1118) |
| $R_{work}$ / $R_{free}$ | 0.2069 (0.4244) /0.2556 (0.4580) | 0.2087 (0.2649) /0.2472 (0.3274) | 0.2414 (0.2497) /0.2913 (0.3192) | 0.2295 (0.4015) /0.2830 (0.4173) | 0.2079 (0.2107) 0.2528 (0.2899) |
| No. atoms | | | | | |
| Protein | 2228 | 1072 | 910 | 2905 | 6614 |
| Ligand/ion | n/a | 2 | 11 | 3 | n/a |
| Water | 92 | 46 | 33 | 19 | 5 |
| *B*-factors | | | | | |
| Protein | 51 | 35 | 40 | 73 | 76 |
| Ligand/ion | n/a | 25 | 57 | 87 | 40 |
| Water | 52 | 41 | 43 | 64 | |
| R.m.s. deviations | | | | | |
| Bond lengths (Å) | 0.008 | 0.010 | 0.006 | 0.002 | 0.002 |
| Bond angles (°) | 0.920 | 0.960 | 0.760 | 0.463 | 0.420 |

*Single Crystal used for each data collection. *Values in parentheses are for highest-resolution shell.

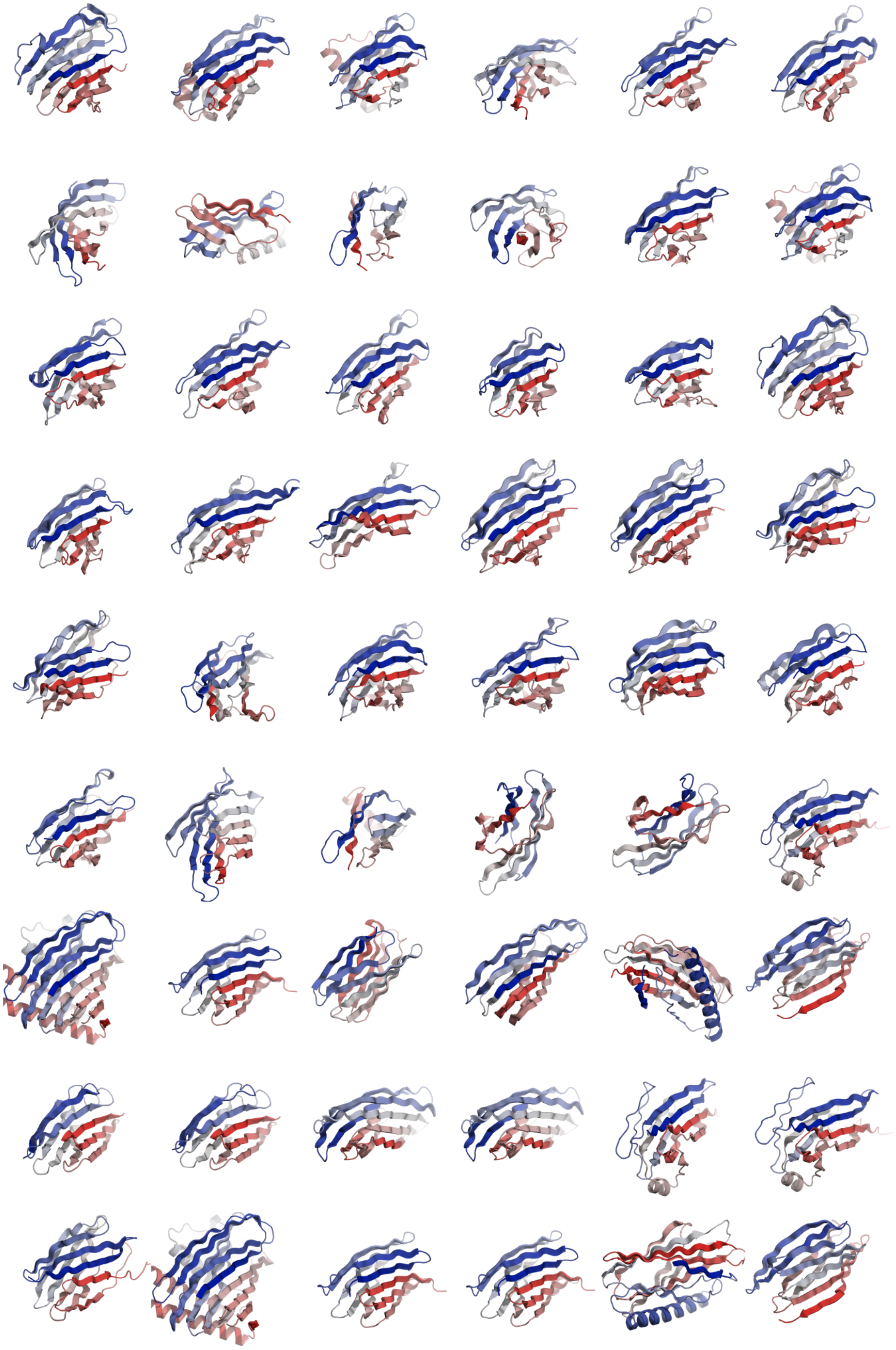

**Fig. S1. Overview of all experimentally confirmed DIPS** (DIP001 - DIP0054) models from left to right, top to bottom, colored from N-terminus (blue) to C-terminus (red).

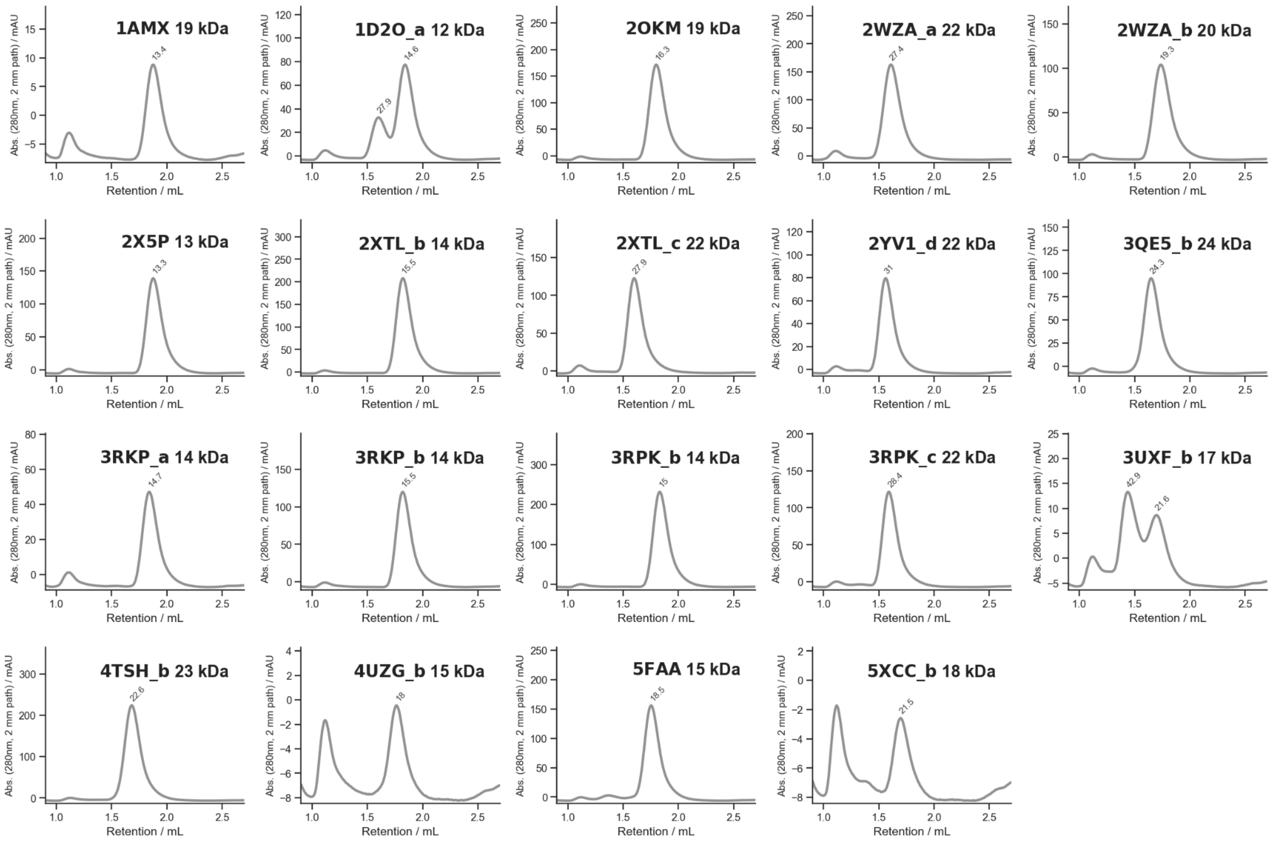

**Fig. S2. SEC traces of native isopeptide bond forming domains (N=19)** used as template, identified by their PDB id and subdomain if the pdb contains multiple isopeptide bond forming domains, for structural models see supporting data or github repository. These are the data for the yield calculator in Fig. 1E. SEC traces recorded on a Cytiva Superdex 75 Increase 5/150 GL in PBS, absorbance at 280 nm, path length annotated), approximate molecular weight extracted from column calibration annotated next to each peak in kDa.

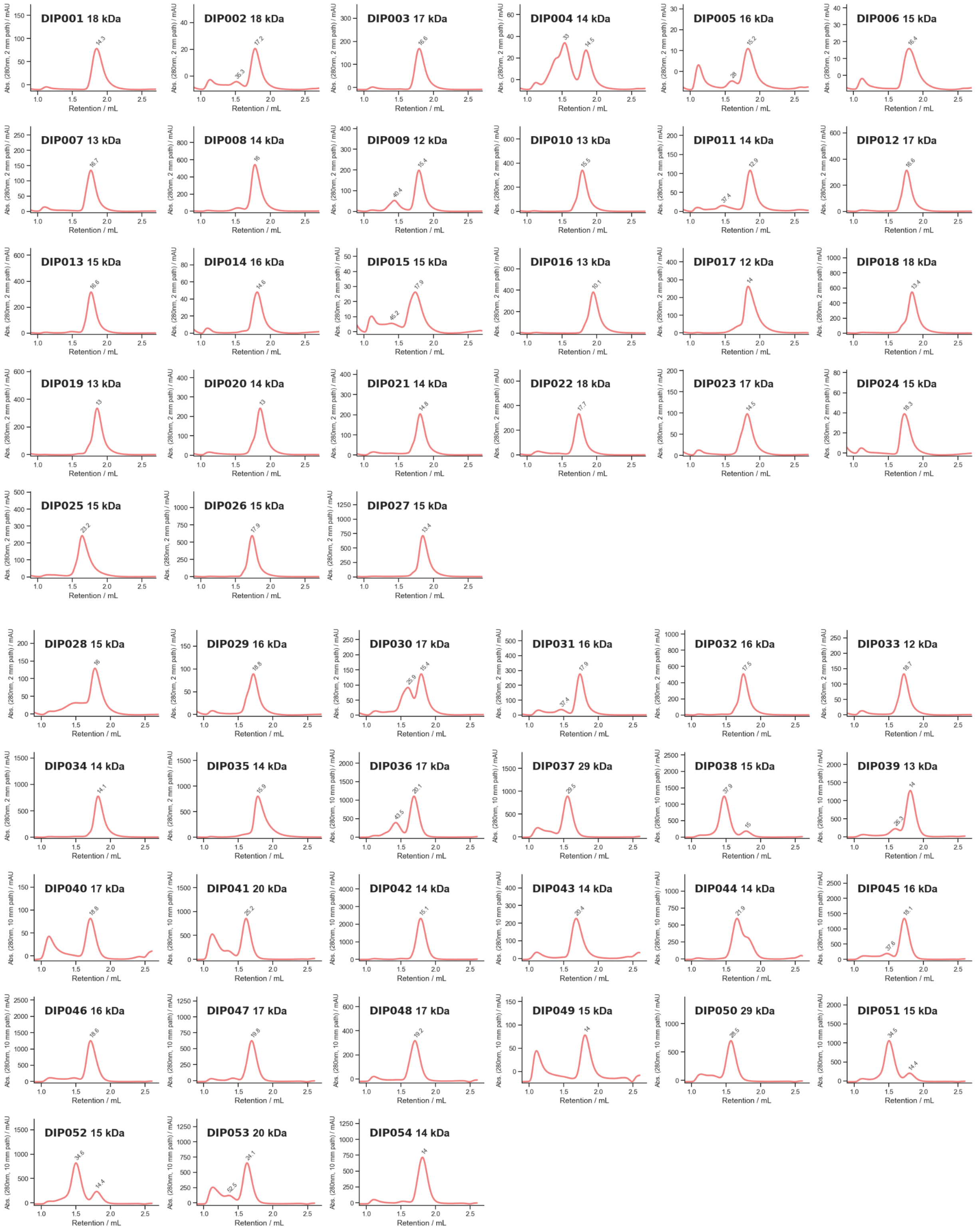

**Fig. S3. SEC traces of all DIPS.** Here shown on a Cytiva Superdex 75 Increase 5/150 GL (approx. 3 mL bed volume) at 0.65 mL / min flow speed in PBS. Absorbance at 280 nm is shown for all designs, though the path length differs between measurements changing the absolute mAU values. SEC traces recorded on a Cytiva Superdex 75 Increase 5/150 GL in PBS, absorbance at 280 nm, path length annotated), approximate molecular weight extracted from column calibration annotated next to each peak in kDa.

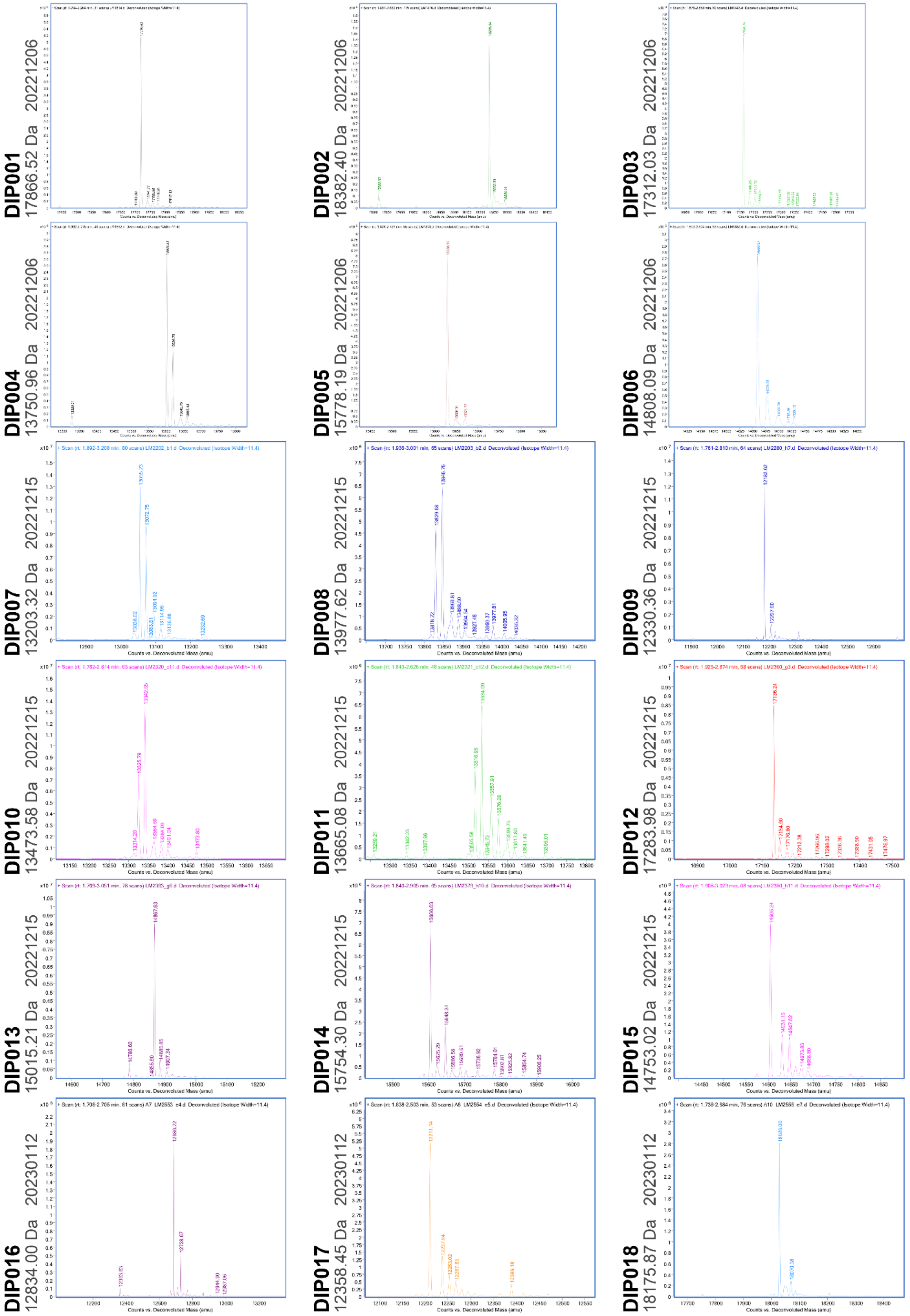

**Fig. S4. Deconvoluted mass spectra of DIPS (DIP001-DIP018).** With the expected masses for full constructs annotated, clearly identifiable peaks of the spectra, though not always the main peak when the isopeptide bond is only formed in a fraction of the proteins analyzed, show a consistent mass loss of around 17 Da when accounting for N-terminal Methionine loss due to processing in *E. Coli* loss.

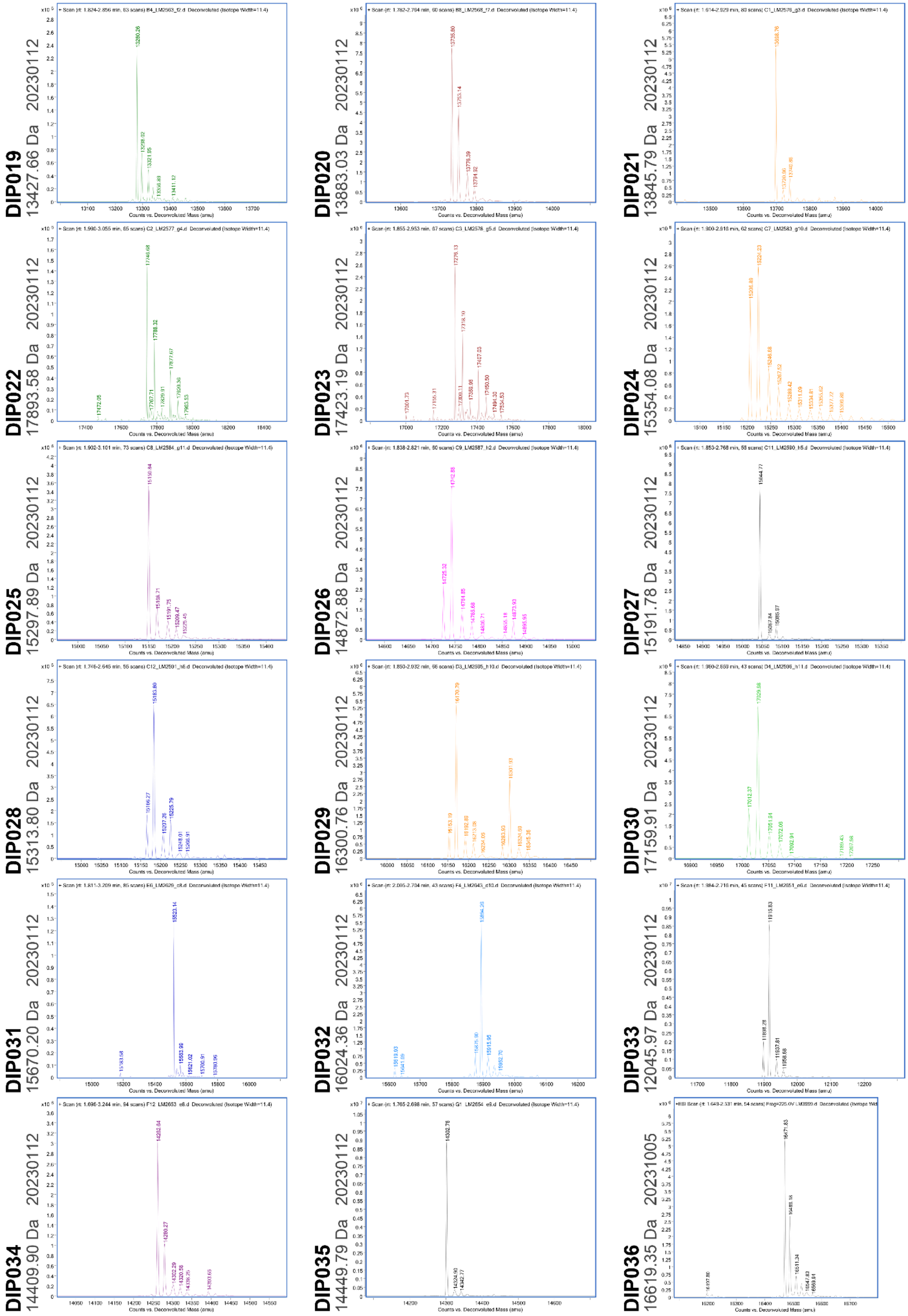

**Fig. S4. continued (DIP019 - DIP036).**

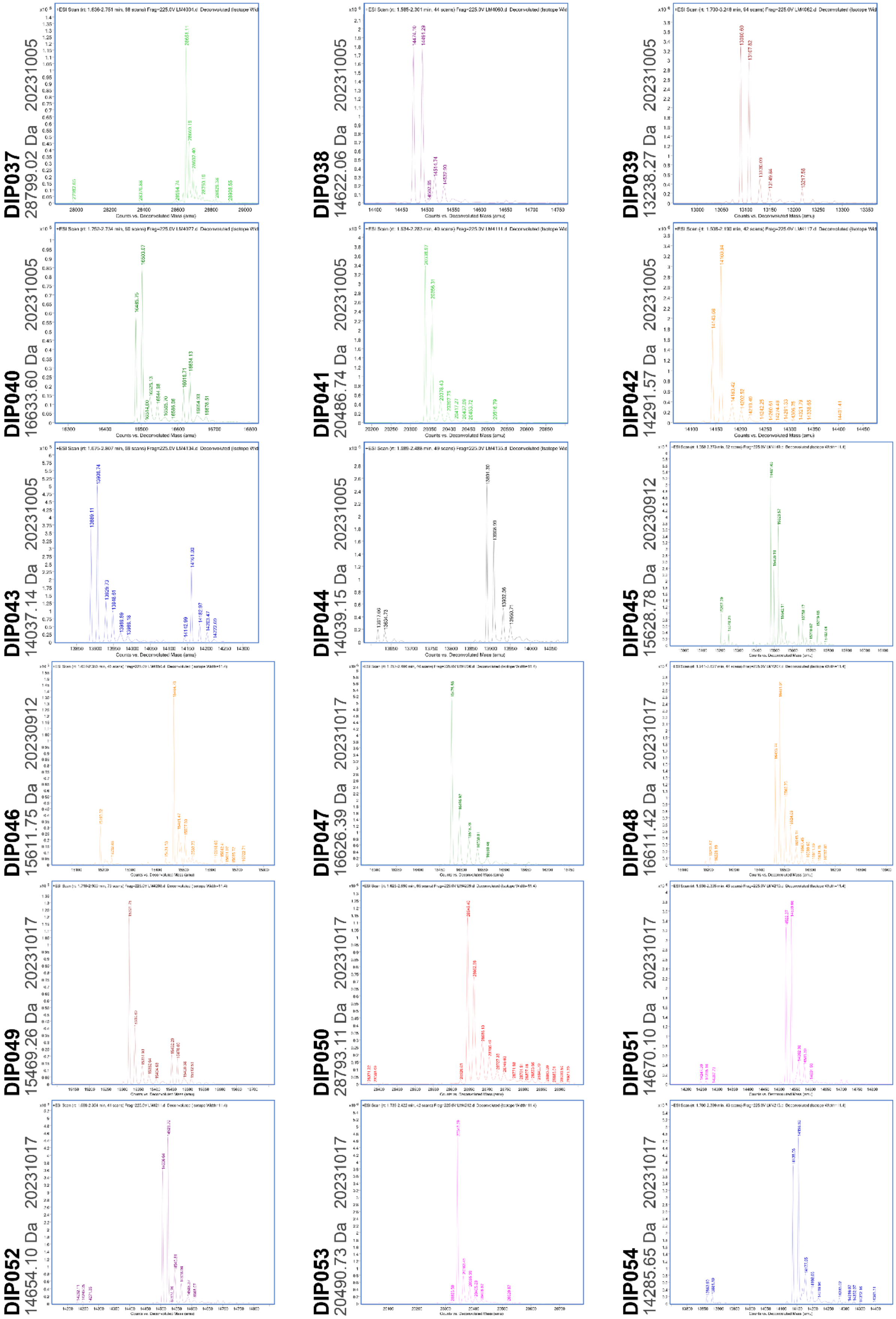

**Fig. S4 continued (DIP037 - DIP054).**

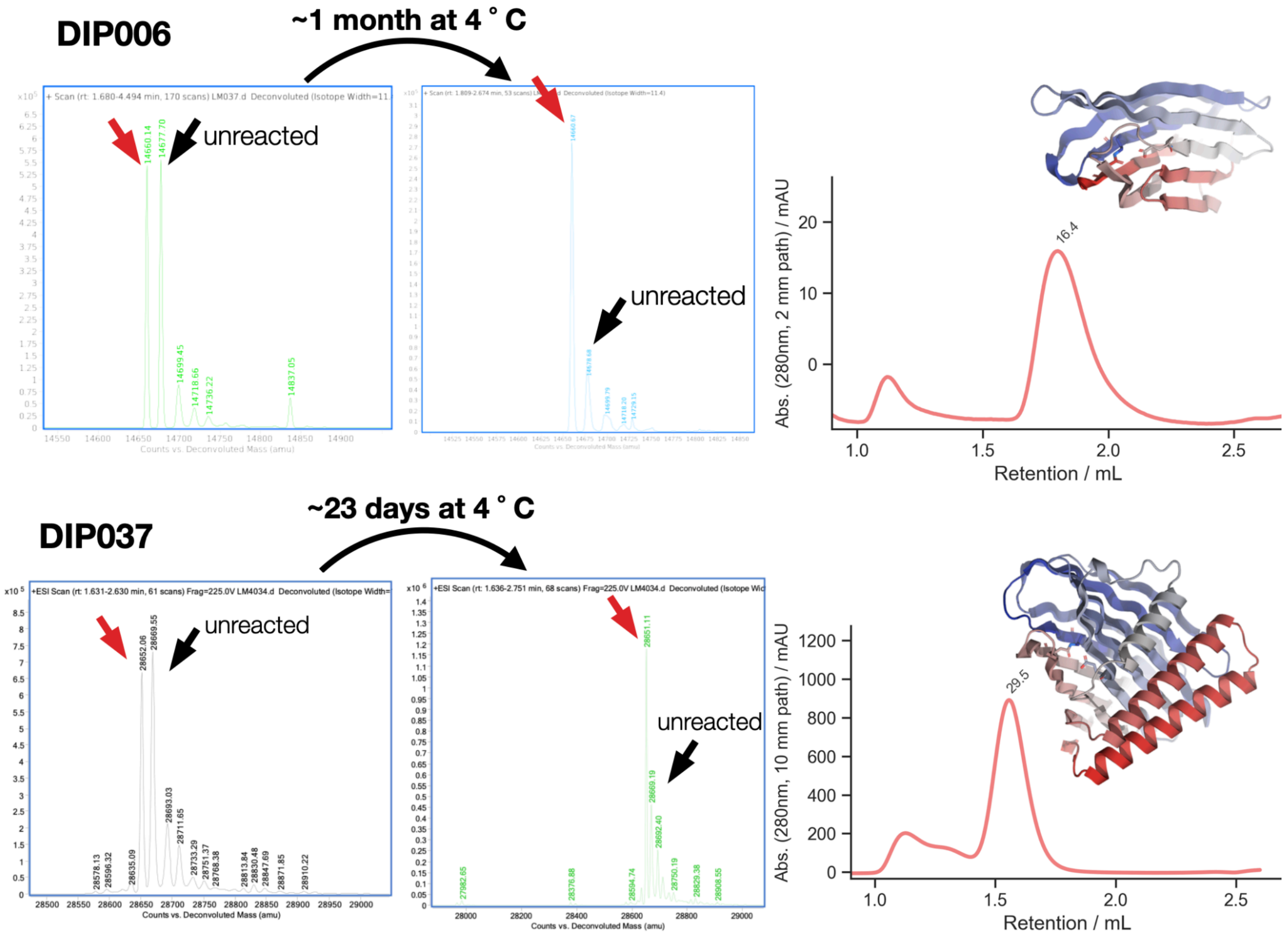

**Fig. S5. Slow forming isopeptide bonds react over weeks**. Here shown for DIP006 and DIP037 SEC on Cytiva S75 5/150 shown with calibrated masses in kDa annotated for each peak on the right, with inset of design models, reacting side chains are highlighted as sticks. Left: Ratio of reacted to unreacted peaks (black arrow) in the deconvoluted mass spectra shift towards the product of an isopeptide bond containing protein fraction with a 17 Da mass loss (red arrow) after accounting for N-terminal Methionine loss, here after over 3 weeks, noticeably the isopeptide bond linked product peak of lighter mass increases.

**A**

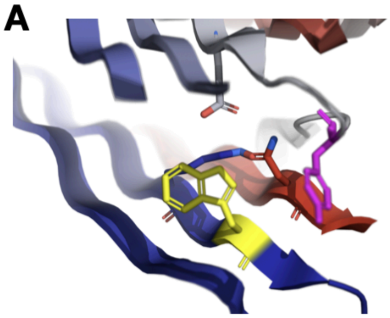

| name | expected mass | LC-MS | Δm | corr-ected for MET | Iso-peptide |
|---|---|---|---|---|---|
| DIP002 | 18382.4 | 18234 | 148.4 | 17.4 | yes |
| DIP002_KO_K7A | 18325.31 | 18194 | 131.31 | 0.31 | no |
| DIP002_KO_E64A | 18324.36 | 18193 | 131.36 | 0.36 | no |
| DIP002_KO_N141A | 18339.38 | 18208 | 131.38 | 0.38 | no |
| DIP002_W9A | 18267.27 | 18136 | 131.27 | 0.27 | no |
| DIP002_Y69A | 18290.31 | 18142 | 148.31 | 17.31 | yes |

**B**

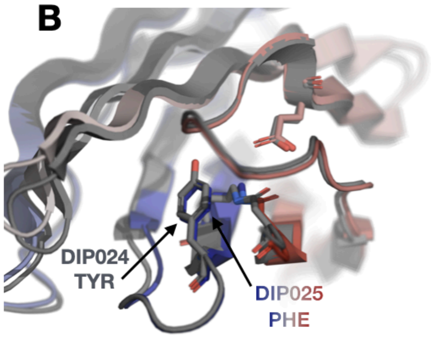

TM-align DIP024 / DIP025
TM-score = 0.97, RMSD 0.74 Å

| name | expected mass | LC-MS | Δm | corr-ected for MET loss | Iso-peptide |
|---|---|---|---|---|---|
| DIP024 main peak | 15354.28 | 15224.23 | 130.05 | -0.95 | no |
| **DIP024 side peak** | 15354.28 | 15206.89 | 147.39 | 16.39 | yes |
| **DIP025 main peak** | 15298.09 | 15150.84 | 147.25 | 16.25 | Yes |

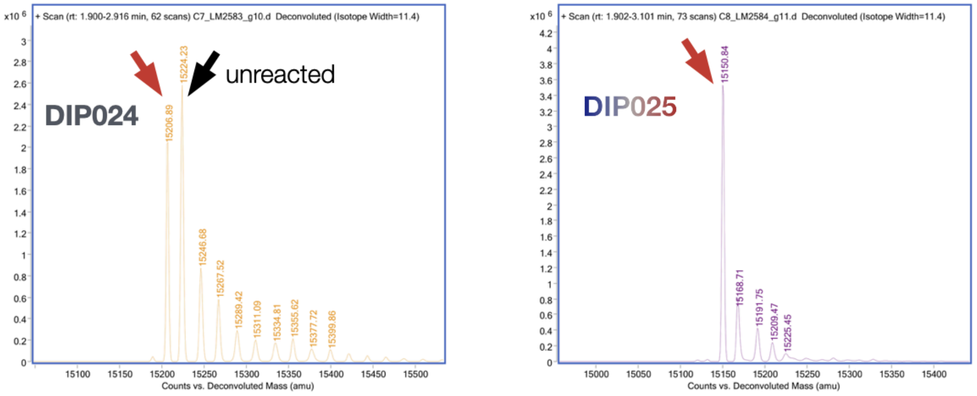

**Fig. S6. Local environment steers isopeptide bond formation rate.**
(A) For DIP002 mutation any of the reactive triad residues (LYS, GLU, or ASN) leads to a loss of isopeptide bond formation, no mass loss of leaving $NH_3$ detected in LC-MS. The same loss of isopeptide bond formation is also observed when the closing hydrophobic TRP (shown in yellow) following the reactive Lysine is mutated to Alanine, highlighting the importance of a hydrophobic environment for the reaction. Finally a conserved TYR (purple) on a closing loop often found in the native structures, can be mutated to ALA without affecting isopeptide bond formation. (B) DIP024 forms an isopeptide bond only partially, the main peak showing no expected $NH_3$ loss, a smaller peak shows that a fraction of DIP024 has formed the bond. DIP025 has 71% sequence identity to DIP024, with a virtually identical protein core, the TM-align backbone RMSD between both design models is < 1 Å. However, DIP025 has a clear difference: a more hydrophobic PHE instead of DIP024 TYR closing over the isopeptide bond, likely this explains why DIP025 forms the isopeptide bond with quantitative yield.

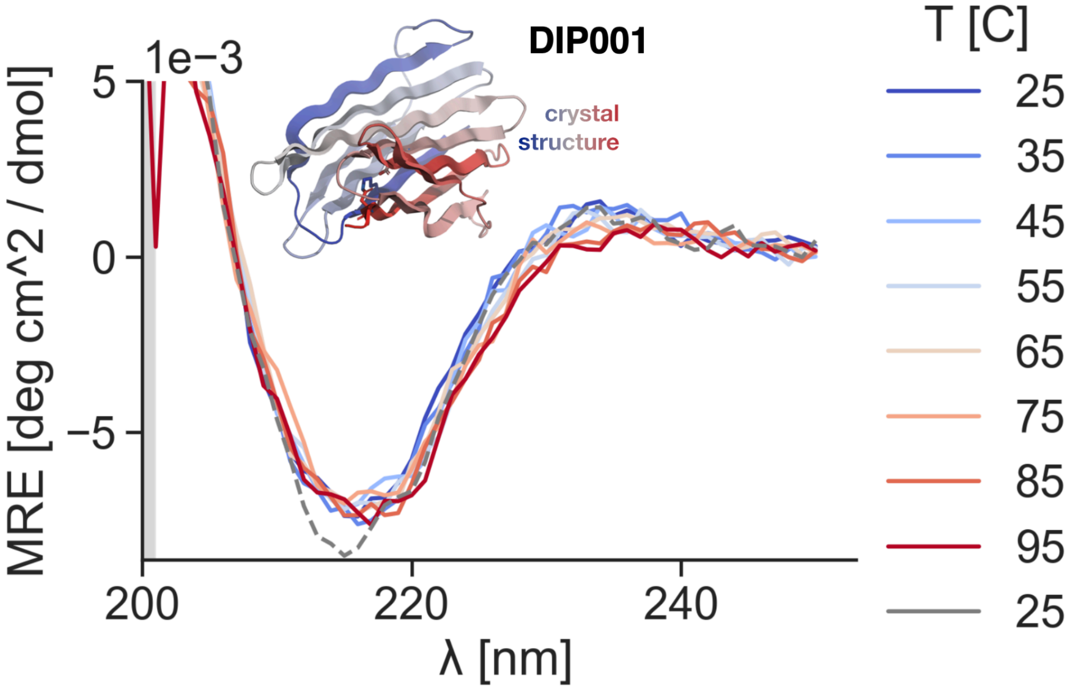

**Fig. S7. Circular Dichroism spectra of DIP001** with its crystal structure shown as inset. Heated from 25 to 95˚C at 0.2 mg/mL concentration in PBS (pH 7.4). The gray dashed line marks the final spectrum recorded after cooling down from 95 ˚C. Spectra show a characteristic dip for beta sheets at around 218 nm, and the spectra do not change between temperatures indicating the protein is stable up 95 ˚C.

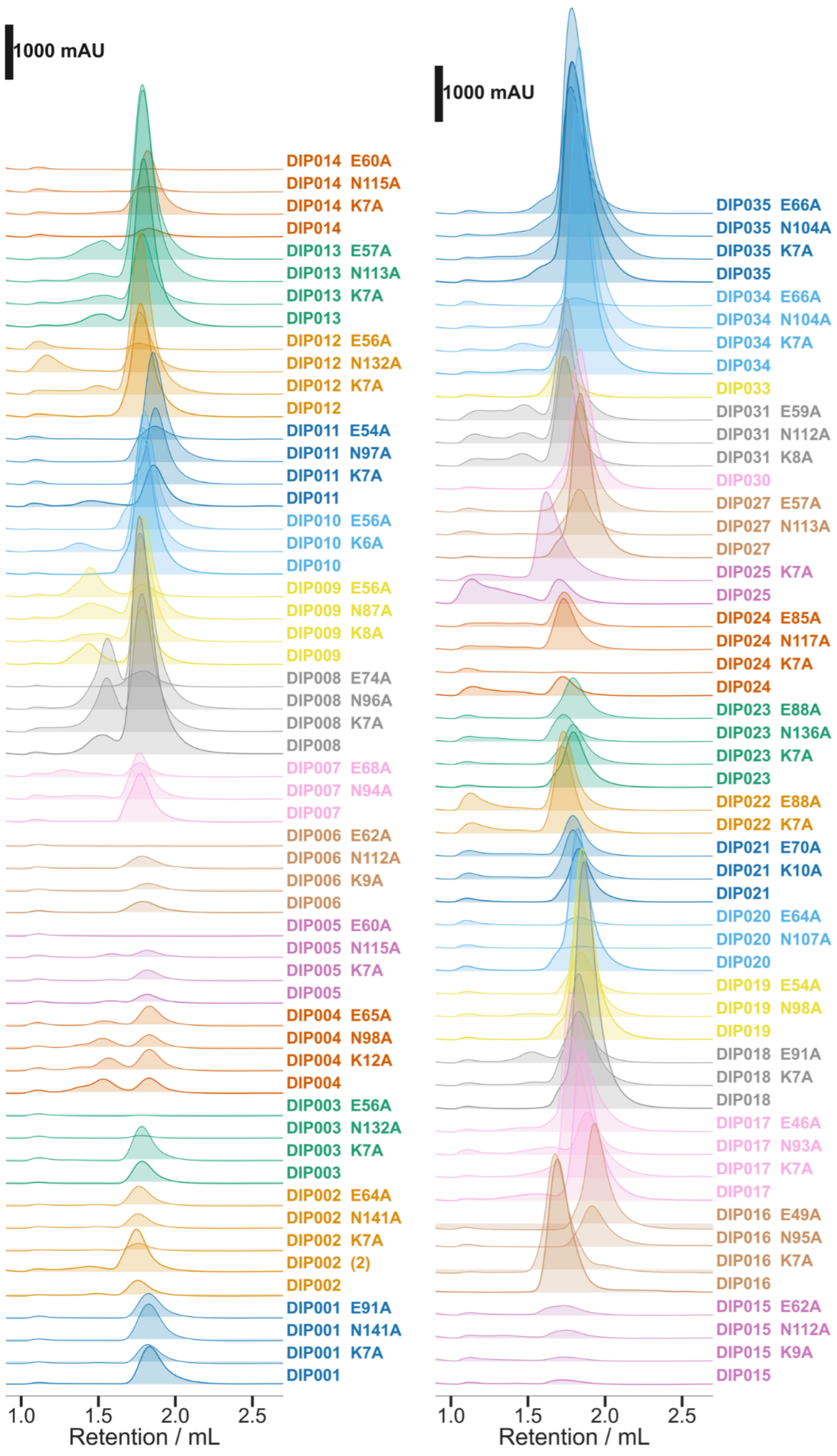

**Fig. S8. Active site Alanine mutations in DIPS often retain solubility for DIP001 to DIP035.** Mutations to ALA of one of each of the active site residues, LYS, catalytic GLU, ASN in designs retains their solubility and often keeps their SEC profiles unchanged. Isopeptide bond formation is not observed for any active site residue to ALA mutant. SEC traces recorded on a Cytiva Superdex 75 Increase 5/150 GL in PBS, absorbance at 280 nm adjusted to 10 mm path length).

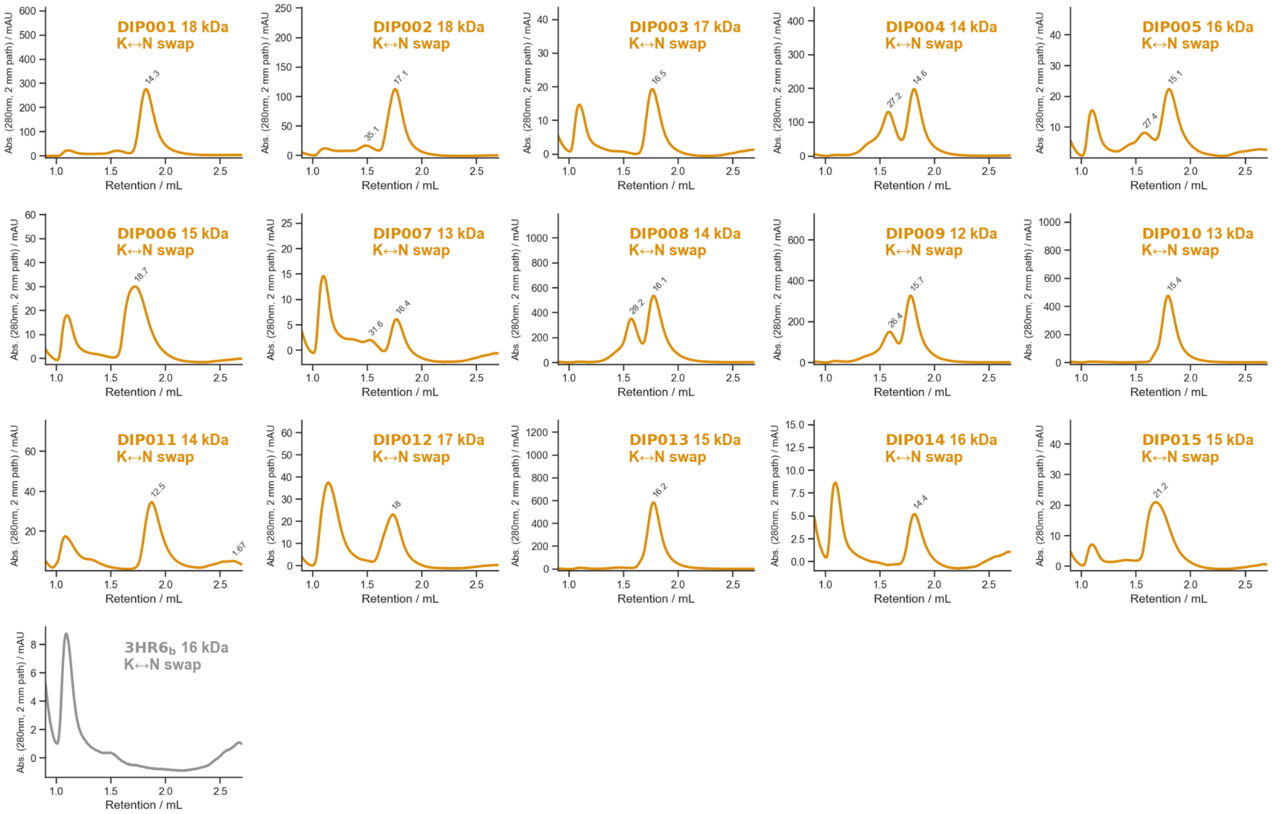

**Fig. S9. Swapping LYS and ASN position in DIPS and one native can produce soluble protein but no isopeptide bond formation.** In these designs the positions of the reacting LYS and ASN of the reactive triad were swapped, remarkably this inhibits isopeptide bond formation completely, but can still produce soluble, monodisperse protein, for example DIP001, DIP002, DIP013. When this swap is applied to a native protein 3HR6_b, no soluble yield can be detected. SEC traces recorded on a Cytiva Superdex 75 Increase 5/150 GL in PBS, absorbance at 280 nm, path length annotated), approximate molecular weight extracted from column calibration annotated next to each peak in kDa.

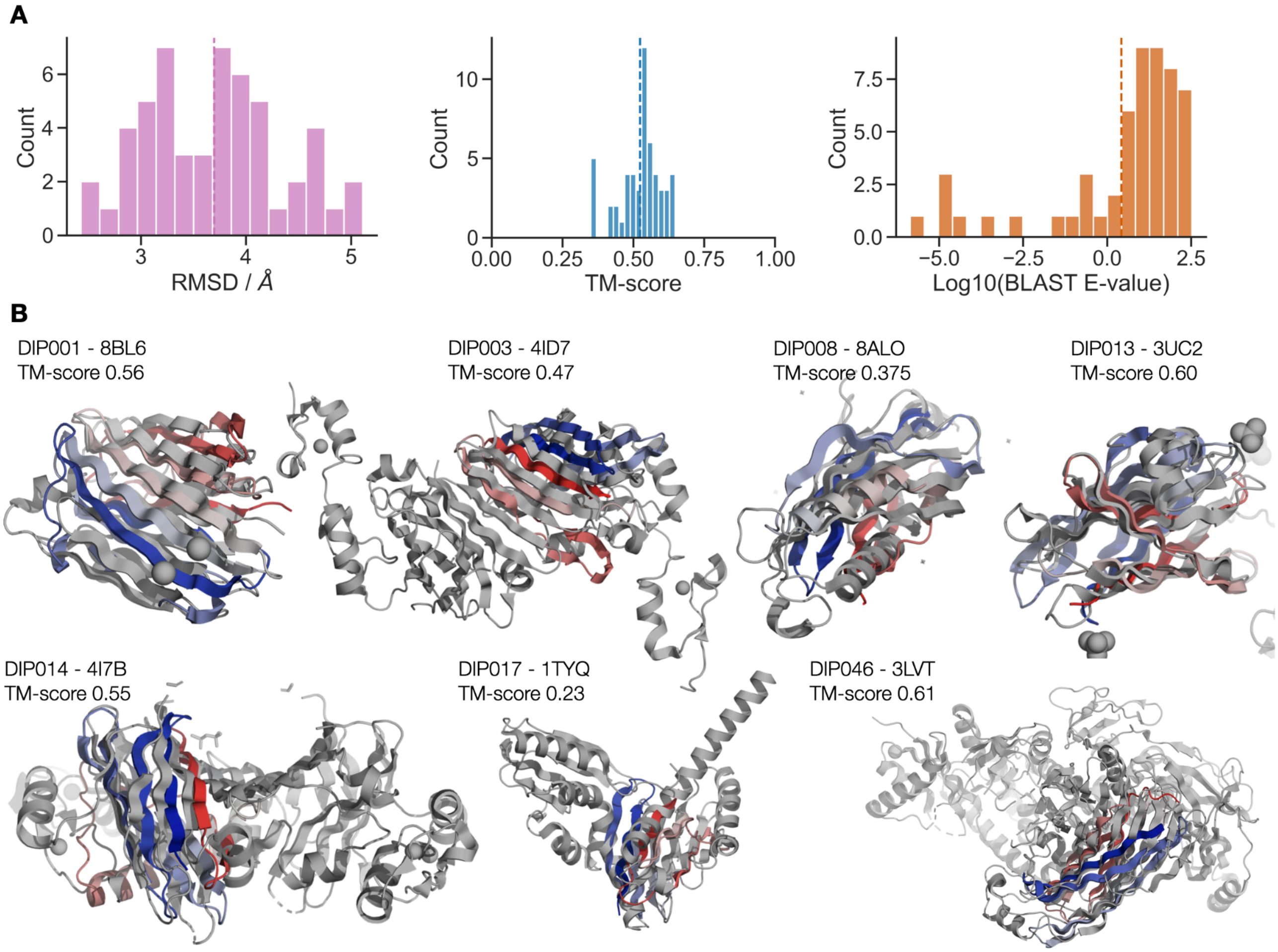

**Fig. S10. Novelty of designs.** For all confirmed isopeptide bond forming designs (N=54), novelty was investigated using: (A) Foldseek Tm-align search against the PDB100. For the respective best PDB structure from the search, histograms show RMSD and TM-score, as calculated by a local TM-align run against that structure, vertical line marks the mean (mean 3.7 A RMSD, mean 0.52 TMscore). (B) Protein BLAST search for each design with lowest E-value result, vertical line marks mean (0.42) (C) Alignment of top hits from the PDB against the crystallized designs (DIP001, DIP003, DIP013, DIP014, DIP046) with PDB IDs of a TM-align against the pdbs from the foldseek search with the highest foldseek TM-score.

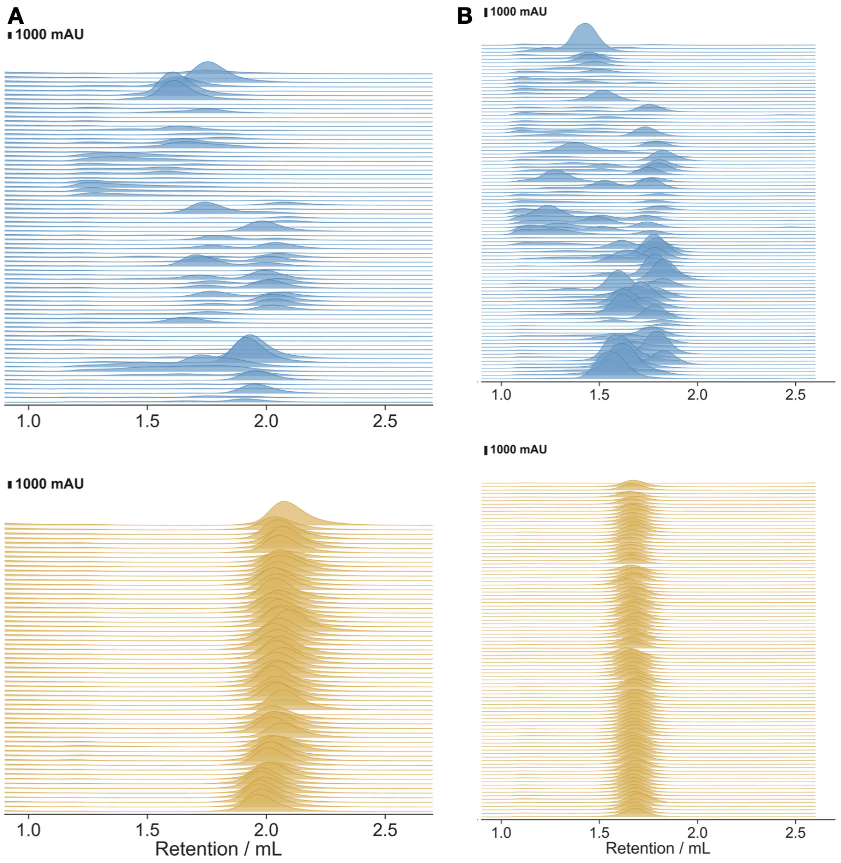

**Fig. S11. Tag Catcher splits SEC traces.** In both design campaigns GG29 (A) and GG32 (B). Noticeably the DIPcatcher domains in blue have worse yield and monodispersity, if they are soluble at all. The peptide DIPtags fused to protein GB1 (A) o FIVAR (B) via a short GS linker have much better yields, monodispersity, and consistent elution volume. SEC traces recorded on a Cytiva Superdex 75 Increase 5/150 GL in PBS, absorbance at 280 nm adjusted to 10 mm path length)

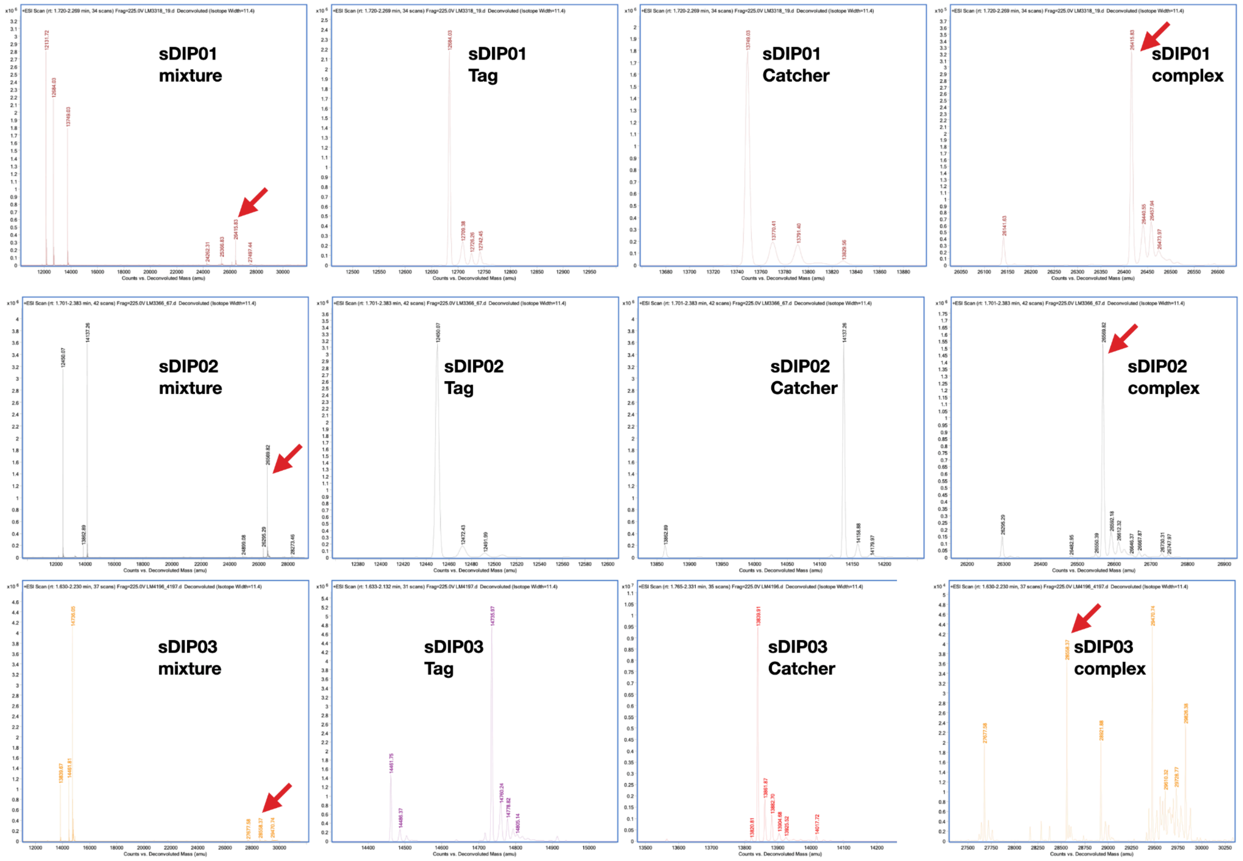

| name | expected mass | LC-MS | Δm | corr-ected for MET loss | Iso-peptide |
|---|---|---|---|---|---|
| sDIP01_Catcher | 13879.37 | 13749.03 | -130.34 | 0.66 | - |
| sDIP01_Tag | 12814.75 | 12684.03 | -130.72 | 0.28 | - |
| **sDIP01_complex** | **26694.12** | **26415.83** | **-278.29** | **-16.29** | Yes |
| sDIP02_Catcher | 14267.57 | 14137.26 | -130.31 | 0.69 | - |
| sDIP02_Tag | 12580.58 | 12450.07 | -130.51 | 0.49 | - |
| **sDIP02_complex** | **26848.15** | **26569.82** | **-278.33** | **-16.33** | Yes |
| sDIP03_Catcher | 13970.33 | 13839.91 | -130.42 | 0.58 | - |
| sDIP03_Tag | 14866.48 | 14735.97 | -130.51 | 0.49 | - |
| **sDIP03_complex** | **28836.81** | **28558.37** | **-278.44** | **-16.44** | Yes |

**Fig. S12. LC-MS detection of isopeptide crosslink for split DIPS (sDIPS01-03).** SplitDIPS sDIP01, sDIP02, and sDIP03 cognate Tag and Catcher complexes were run on LC-MS after at least 48h incubation times (in PBS at equimolar concentration of 10 µM each). The deconvoluted mass spectra are displayed with zoomed in peaks for the individual peaks that correspond to the masses of the DIPcatcher and DIPtag (here fused to FIVAR or SUMO as solubility tags). The summary table shows the formation of an isopeptide bond between each Tag-Catcher pair as indicated by the presence of a peak that is the added mass of Tag and Catcher that, after accounting for N-terminal MET loss off both proteins, shows a mass loss of approx. 17 Da, the $NH_3$ lost on isopeptide bond formation, indicating that they are covalently linked via an isopeptide bond.

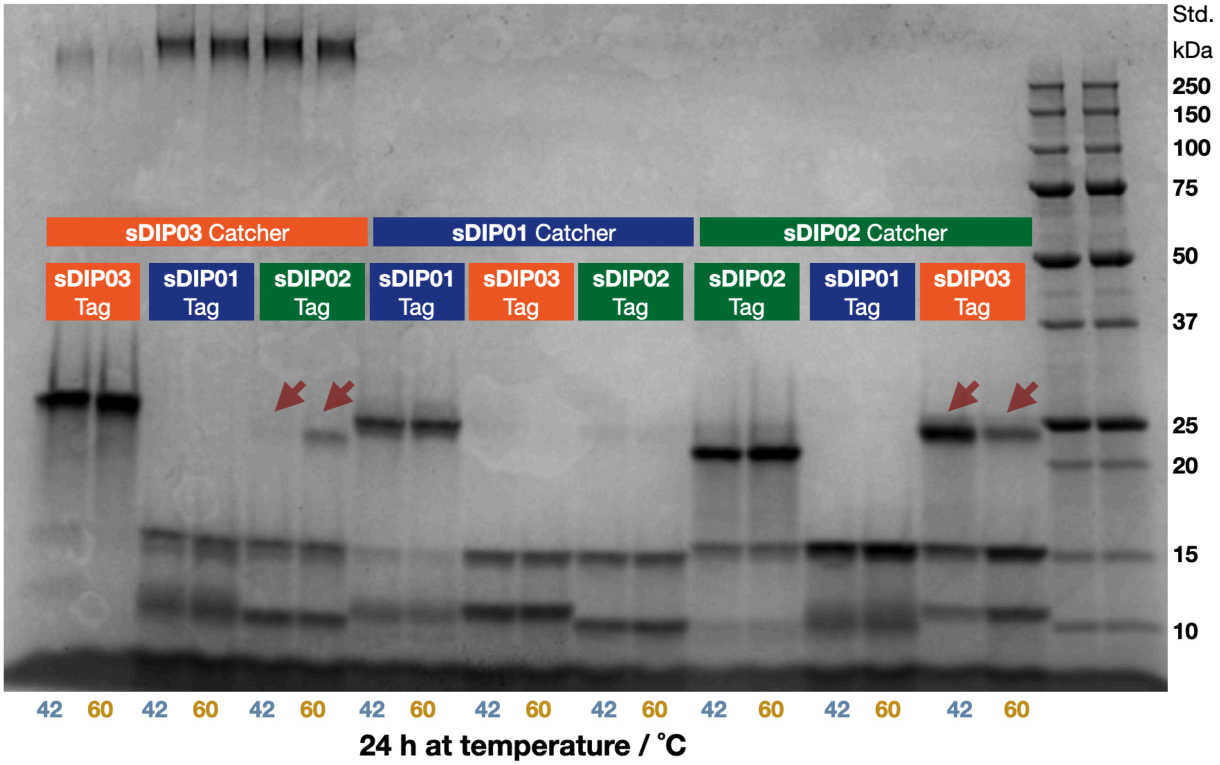

**Fig. S13. sDIP crossreactivity testing validated sDIP Catcher domains.** (marked in matching colors at the top, each approx. 14 kDa) against all sDIPtags (here fused to a heat stable version of protein A as mass tag approx. 10 kDa), here incubated for 24 h at either 42 or 60 ˚C (marked at the bottom) in PBS with a final concentration of 10 μM for each binding partner in the reaction and then run on a denaturing SDS-PAGE gel. Cross reactivity covalent product formation is indicated by formation of approx 25 kDa molecular weight band on an SDS-PAGE with Coomassie blue staining between tag and catcher, indicated with red arrows. sDIP02 catcher und sDIP03 tag, as well as, but less strongly, sDIP03 catcher and sDIP02 tag, cross react in an undesired way.

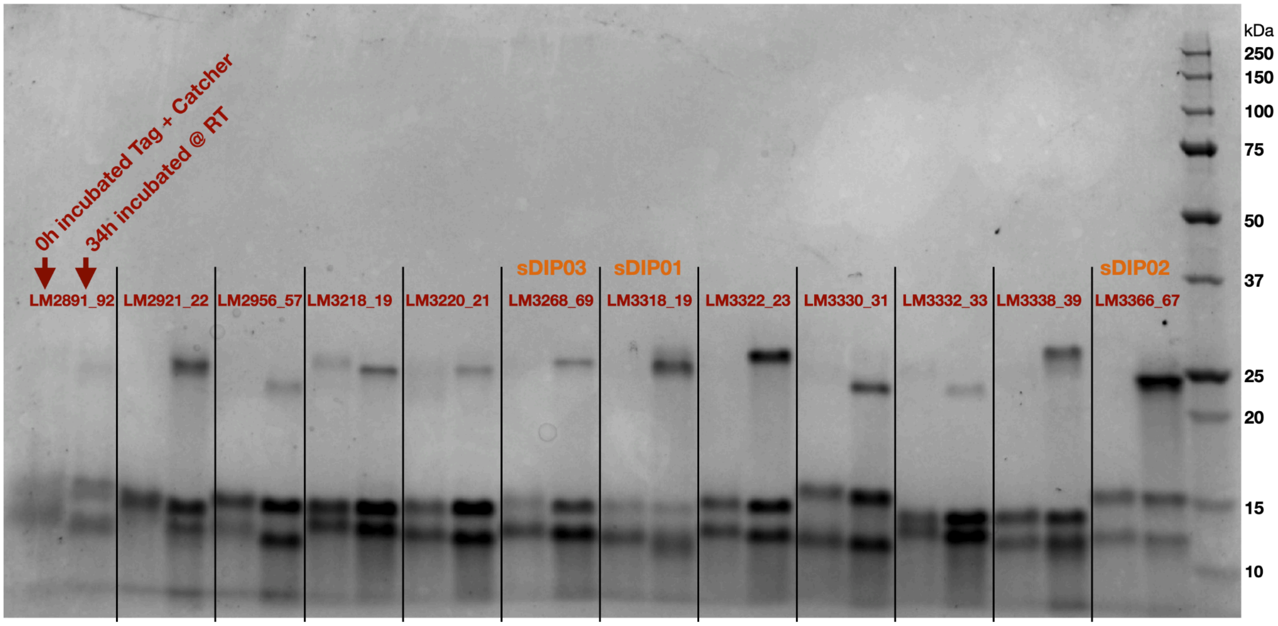

**Fig. S14. Other sDIP candidates.** that were ultimately not developed further. Here shown at 10 μM of each binding partner in PBS (pH 7.4) incubated at room temperature (RT) either for 0h or 34h, covalent product formation linked by an isopeptide bond can be detected for all, albeit at different ratios. The chosen sDIPS01-03 are marked. Other designs are given as e.g. LM2921_22, meaning designs LM2921 Catcher + LM2922 Tag (sequences provided in Supplementary Data).

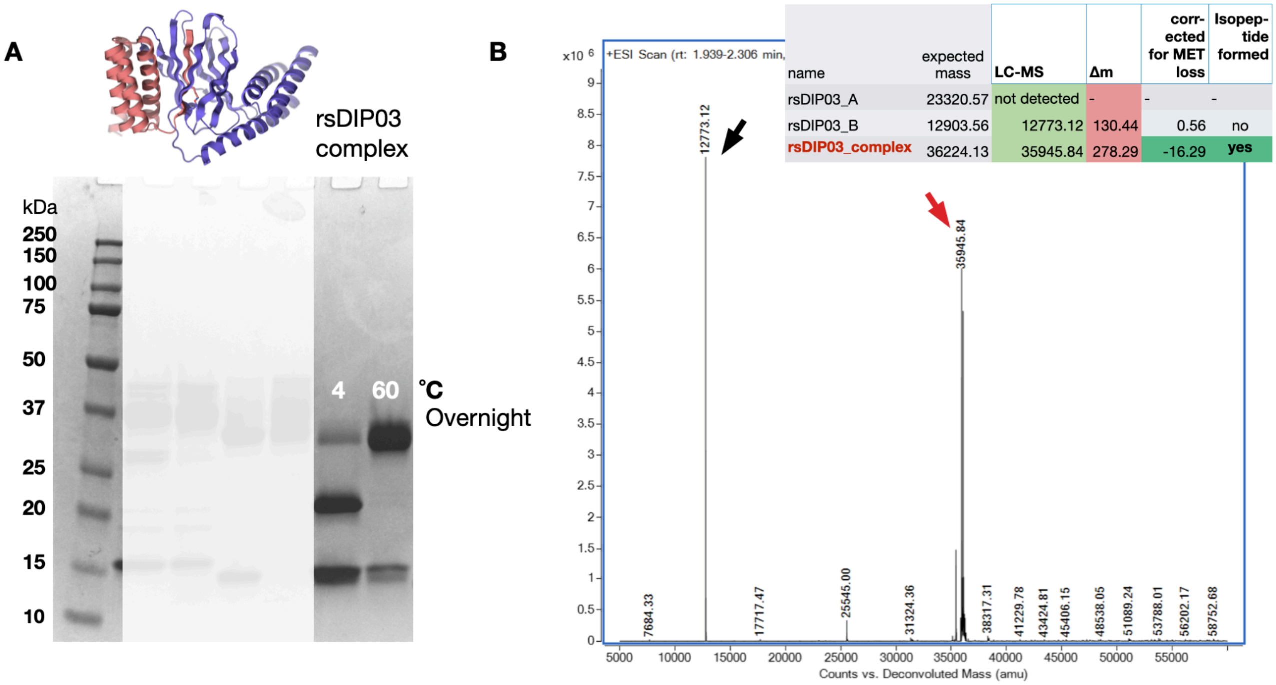

**Fig. S15. rsDIP03 isopeptide bond formation validation.** (A) Rigid split DIP rsDIP03, design model displayed, reacts to form a covalent product after overnight incubation at both 4 and 60 ˚C, although only at the higher temperature the entirety of the larger rsDIP03_A (around 23 kDa) has reacted with the smaller rsDIP03_B subunit (approx. 13 kDa) to form an isopeptide linked product (approx. mass 36 kDa), here indicated by SDS-PAGE stained with Coomassie. (B) The isopeptide linked product can be detected with its expected mass on LC-MS. When correcting for MET loss on each subunit, a mass loss of approx 16 Da of the product is observed, close enough to the 17 Da expected for $NH_3$ loss upon isopeptide bond formation is detected in the deconvoluted LC-MS spectra.
